# Ant And Mite Diversity Drives Toxin Variation In The Little Devil Poison Frog

**DOI:** 10.1101/031849

**Authors:** Jenna R. McGugan, Gary D. Byrd, Alexandre B. Roland, Stephanie N. Caty, Nisha Kabir, Elicio E. Tapia, Sunia A. Trauger, Luis A. Coloma, Lauren A. O’Connell

## Abstract

Poison frogs sequester chemical defenses from arthropod prey, although the details of how arthropod diversity contributes to variation in poison frog toxins remains unclear. We characterized skin alkaloid profiles in the Little Devil frog, *Oophaga sylvatica* (Dendrobatidae), across three populations in northwestern Ecuador. Using gas chromatography mass spectrometry, we identified histrionicotoxins, 3,5- and 5,8-disubstituted indolizidines, decahydroquinolines, and lehmizidines as the primary alkaloid toxins in these *O. sylvatica* populations. Frog skin alkaloid composition varied along a latitudinal gradient across populations in a principal component analysis. We also characterized diversity in arthropods isolated from frog stomach contents and confirmed *O. sylvatica* specialize on ants and mites. To test the hypothesis that poison frog toxin diversity reflects species and chemical diversity in arthropod prey, we (1) used liquid chromatography mass spectrometry to chemically profile consumed ants and mites, and (2) used sequencing of cytochrome oxidase 1 to identify individual prey specimens. We show that chemical profiles of consumed ants and mites cluster by frog population, suggesting different frog populations have access to chemically distinct prey. We identified 45 ants and 9 mites isolated from frog stomachs, finding several undescribed species. Finally, by comparing chemical profiles of frog skin and isolated prey items, we were able to trace the arthropod source of four poison frog alkaloids, including 3,5- and 5,8-disubstituted indolizidines and a lehmizidine alkaloid. Together, our data shows the diversity of alkaloid toxins found in *O. sylvatica* can be traced to chemical diversity in arthropod prey.

## INTRODUCTION

Many organisms have evolved sophisticated chemical defenses to deter predators. Some defended organisms produce their own chemical defenses while others sequester toxins from external sources (Casewell et al. 2013; Olivera et al. 1985; Saporito et al. 2012; Saporito et al. 2009). Toxin sequestration is best understood in invertebrates, where examples of plant to arthropod toxin transfer has been described in great detail (Heckel 2014; Opitz and Müller 2009). Compared to invertebrates, far less is known about how vertebrate species acquire chemical defenses from external sources. An example of toxin sequestration among vertebrates is poison frogs, which accumulate alkaloid toxins from arthropod prey (Daly et al. 1994a; Daly et al. 1994b; Hantak et al. 2013; Saporito et al. 2009). Although the dietary basis of poison frog toxicity is well established, the relationship between arthropod prey diversity and frog toxin diversity is not fully understood (Santos et al. 2015; Saporito et al. 2007b; Saporito et al. 2012). Here we characterize the toxins found on the Little Devil frog *(Oophaga sylvatica)* and their arthropod prey and trace the dietary source of specific alkaloids to the ants and mites they ingest.

The term "poison frog” refers to anurans that carry alkaloid toxins in their skin and include several families in South America such as dendrobatids (Dendrobatidae) and bufonids *(Melanophryniscus)*, as well as mantellids *(Mantella)* from Madagascar, myobatrachids *(Pseudophryne)* from Australia, and Eleutherodactylids from Cuba (Daly 1995; Daly and Spande 1986; Rodríguez et al. 2011). Many poison frogs are diurnal and brightly colored, presumably serving as an aposematic signal to predators (Darst and Cummings 2006; Darst et al. 2005; Santos et al. 2003; Santos et al. 2015). Many of these chemically defended frogs carry small molecule alkaloid toxins in granular glands on their skin (Neuwirth et al. 1979). Decades of work in poison frog chemical ecology has identified over 800 alkaloids organized into over 22 structural classes (Daly et al. 2005; Saporito et al. 2012), although the structure of many unclassified frog alkaloids remains to be determined. Some of these alkaloids are highly toxic to vertebrates, such as batrachotoxin and pumiliotoxin, while many others like histrionicotoxins and decahydroquinolines are noxious or bitter tasting (Santos et al. 2015). Most of the work on poison frog chemical ecology has focused on the Strawberry poison frog *(Oophaga pumilio)*. Over 250 alkaloids have been identified in *O. pumilio* with variation both between and within populations (Saporito et al. 2006; Saporito et al. 2007a; Saporito et al. 2010). However, the ecological drivers and genetic contributions to variation in chemical profiles are still not understood. More integrative and comparative work is needed in order to comprehend how environmental and genetic variables contribute to the diverse repertoires of defensive chemicals in poison frogs.

Initial reports on poison frog alkaloid toxins suggested frogs synthesized these chemicals. Once poison frog captive colonies were established, however, researchers determined the frogs had lost their toxicity when fed a diet of fruit flies. These observations led to the dietary hypothesis for poison frog toxicity (Daly et al. 1994a; Daly et al. 1994b), and since then ecological studies have demonstrated that poison frogs ingest mainly ants and mites (Caldwell 1996; Donnelly 1991; Toft 1980), that this dietary specialization correlates with toxicity (Darst et al. 2005), and that the sequestration of alkaloid toxins from arthropod prey has independently evolved at least four times in the Dendrobatidae family (Santos et al. 2003). Presumably, these toxic species have evolved sequestration mechanisms that allow the binding and transport of toxins from the gut to the skin, although this pathway has not yet been identified (Santos et al. 2015; Saporito et al. 2012). Nevertheless, as poison frogs sequester their toxins from arthropod prey, it has been proposed that the toxin diversity observed within and between species is reflective of arthropod diversity in the tropics (Saporito et al. 2012; Saporito et al. 2009). This implies that sequestration mechanisms in poison frogs must be broad enough to sequester a range of chemicals that take advantage of local arthropod chemistry. Understanding how poison frogs acquire toxins and how differences in diet influence toxin variability and the maintenance of aposematism remains a challenge.

Most research efforts to identify the dietary source of frog alkaloids have focused on arthropods collected from leaf litter. Ants have been identified as a source of many poison frog alkaloids including pumiliotoxins, histrionicotoxins, 5,8 and 3,5-disubstituted indolizidines, decahydroquinolines and pyrrolizidines (Heckel 2014; Jones et al. 1999; Saporito et al. 2004). Despite their small size, mites seem to confer the greatest diversity of alkaloids to poison frogs. Oribatid mites in particular have been explored as the source of many poison frog toxins, including pumiliotoxins and indolizidines (Saporito et al. 2007b; Saporito et al. 2011; Takada et al. 2005). Although there is a clear connection between alkaloid-containing arthropods and poison frog specialization on these ants and mites, little is known about the specific arthropod species poison frogs ingest and the alkaloids these arthropods carry. This is especially the case for mites, whose taxonomy is far understudied compared to ants (Saporito et al. 2015). Although broad leaf litter collections of arthropods and subsequent chemical analyses are useful (Daly et al. 2002; Saporito et al. 2004; Saporito et al. 2009), integrative analyses that include arthropod identification from frog stomachs, chemical analyses of arthropod prey items, and how the arthropod chemical diversity corresponds to poison frog toxin diversity are needed as the next step toward causatively linking poison frog skin toxins to their dietary source.

To determine how arthropod diversity contributes to toxin diversity in a poison frog, here we characterize the alkaloid composition and diet of the Little Devil frog (or Diablito frog, *Oophaga sylvatica)* and trace the dietary source of toxins to ants or mites. We test the hypothesis that ingested ant and mite species differ across frog populations, and that the chemical diversity of ingested prey contributes to corresponding differences in alkaloid variation across frog populations. We first use gas chromatography mass spectrometry (GCMS) to profile the chemical diversity of three Little Devil populations. Then, to test the hypothesis that variation in poison frog toxin profiles reflects the chemical diversity of arthropod prey, we use liquid chromatography mass spectrometry (LCMS) to chemically profile consumed ants and mites. We also identify prey items isolated from frog stomachs using the DNA barcode cytochrome oxidase 1 (CO1) (Meusnier et al. 2008). Finally, we compare the chemical profiles of frogs and their prey items in order to identify either ants or mites as the source of specific alkaloids found in different frog populations.

## METHODS AND MATERIALS

### Field collection

Little Devil *(Oophaga sylvatica)* frogs were collected during the day near the villages of Cristóbal Colón (N=10), Simón Bolívar (N=10) and along the Felfa River (N=12) near the village Montalvo in the northwestern Esmeraldas province of Ecuador in July 2014. Collections and exportation of specimens were done under permits (001-13 IC-FAU-DNB/MA, CITES 17V/S) issued by the Ministerio de Ambiente de Ecuador. Frogs were individually stored in plastic bags with air and vegetation for 3-8 hours. In the evening the same day of capture, frogs were anesthetized with a topical application of 20% benzocaine and euthanized. The dorsal skin (from the back of the head but not including the legs) was isolated and stored in 100% methanol. The stomachs were dissected and their contents were sorted by arthropod type (as genera could not be visually determined) into separate tubes of ants, mites, and other arthropods. Arthropods were stored in 100% methanol at 4°C for a few weeks. To have a nontoxic control group, we also collected five frogs near Otokiki (Province Esmeraldas) and fed them crickets and fruit flies for three months in captivity. The Institutional Animal Care and Use Committee of Harvard University approved all procedures (Protocol 15-02-233). Remaining frog tissues were preserved in 100% ethanol and deposited in the amphibian collection of Centro Jambatu de Investigatión y Conservatión de Anfibios in Quito, Ecuador (CJ 3089-3139). In order to protect the vulnerable *O. sylvatica* populations that are highly targeted by illegal poaching, specific GPS coordinates of frog collection sites can be obtained from the corresponding author.

### Quantification of arthropods in stomach contents

Each arthropod was individually photographed using a Lumenera Infinity 2 camera mounted on an Olympus dissection microscope (SZ40) and assigned a unique seven digit identification number with the first four digits being the voucher specimen number of the frog from which the arthropod was taken and the last three digits being the number assigned to the arthropod in increasing order from which it was removed from the stomach contents. Ant and mite samples from the stomach contents of 5 frogs from each population were selected for alkaloid analysis using liquid chromatography mass spectrometry (LCMS) based on the largest quantity of mites recovered. These samples for LCMS were pooled by arthropod type (ants, mites or other) into 100% methanol until alkaloid extraction. The arthropod samples from the remaining frogs were individually placed in vials of 100% ethanol for later molecular identification by PCR.

Diet was quantified both by quantity of each arthropod type (ants, mites, beetles or "other”) and by volume of each arthropod type to account for the wide range of prey size. To determine volume, the photographs of each arthropod were analyzed using Image J (National Institute of Health, Bethesda, MD) to determine their dimensions. Length measurements were taken from the tip of the mandible and extended to the rearmost point of the arthropod. The width measurement was taken at the midpoint of the arthropod and excluded the extra girth added by appendages. If the prey item was fragmented, the measurements were taken from the nearest identifiable body part. The length and width measurements were used to calculate the volume of each prey item. The equation of a prolate sphere was used for the volume calculation of the ant, beetle, and other arthropods: *V* = (4*π*/3) * (*Length/*2) * *(Width/*2)⌃2. The different body shapes among the mites taken from the stomach contents required the use of three different formulas to most accurately describe their volume and shape. In addition to the equation for a prolate sphere, the equations of a hemi-cylinder (VHC) and a sphere (VS) were also used based on individual mite shape: *VHC* = (4*π*/6) * (*Length*/2) * *(Width/*2)⌃2 and *VS* = *(4*π*/*3) * *(Diameter/*2)⌃3.

### Isolation of alkaloids

A set of samples including frog skins, stomach-isolated ants and stomach-isolated mites from five frogs across populations of Felfa, Simón Bolívar, and Cristóbal Colón were used to characterize alkaloid profiles (45 samples in total). Five additional skin samples from captive frogs (described above) were used as a control. The contents of each sample vial (including arthropods and 100% methanol) were emptied into a sterilized Dounce homogenizer. The empty vial was rinsed with 1 mL of methanol and added to the homogenizer to ensure the full transfer of all materials. As an internal standard, 25 μg of D3-nicotine in methanol (Sigma-Aldrich, St. Louis, MO) was added to each vial. The sample (either frog skin or arthropods) was ground with the piston ten times in the homogenizer before being transferred to a glass vial. The homogenizer and piston were rinsed with 1 mL of methanol that was then also added to the glass vial to collect any alkaloid residue. The equipment was cleaned with a triple rinse of methanol before being used to process another sample. A 200 μL aliquot of sample was removed for later LCMS analysis. The remainder of the frog skin samples was evaporated to dryness under nitrogen gas, reconstituted in 0.5 mL of methanol by vortexing. The samples were transferred to a microcentrifuge tube and spun at 12000 rpm for 10 min. A 200 μL aliquot of the supernatant was transferred to a 0.3 mL glass insert in an amber sample vial for later GCMS analysis. All samples were stored at -20°C until GCMS (frog skin only) or LCMS (frog skin, ants, and mites) analyses.

### Gas chromatography mass spectrometry (GCMS)

GCMS analysis was based on a slight modification of the method reported by (Saporito et al. 2010). Analyses were performed on a Waters Quattro Micro system (Beverly, MA) with an Agilent 6890N GC (Palo Alto, CA). A J&W DB5ms 30 m × 0.25 mm column with a 0.25 μm film thickness (Agilent Technologies, Palo Alto, CA) was used for separations. Helium carrier gas was held constant at 1 mL/min and split injection (split flow ratio of 12) was used with an injector temperature of 280°C. A 2 μL injection of the concentrated and reconstituted methanol extracted samples were used for GCMS analysis. The column temperature gradient began with a 1 min hold at 100°C then increased at 8°C/min to 280°C and was held for 2.5 min. The interface temperature was 280°C. The mass spectrometer was operated in the 70 eV electron ionization mode and scanned continuously in the range m/z 20-616 at a rate of 0.4 sec/scan. Each total ion chromatogram was reviewed and alkaloid responses were characterized from their mass spectra and retention times. The extensive database developed by (Daly et al. 2005) was used to tentatively identify the alkaloids by matching major mass spectrum peaks and relative retention times using D3-nicotine as a reference.

### Liquid chromatography mass spectrometry (LCMS)

LCMS analyses were primarily performed on a Bruker maXis Impact Q-TOF system (Billerica, MA) with an Agilent 1290 LC (Palo Alto, CA). Product ion scans of the ant/mite/skin correlation study were performed on an Agilent 6550 Q-TOF (Palo Alto, CA) configured with the same LC system and operated with similar MS parameters. A reversed-phase LC gradient method was developed using a Phenomenex Gemini C18 3 μm 2.1 × 100 mm column (Torrance, CA). Mobile phase A was composed of water with 0.1% formic acid and mobile phase B was composed of acetonitrile with 0.1% formic acid. The flow rate was 0.2 mL/min. The gradient began with 0% B for one min then increased linearly to 100% B at 15 min and was held until 18 min. LCMS is much more sensitive than GCMS and a 1 μL injection volume of the unconcentrated arthropod samples and a 20-fold diluted (with methanol) frog skin sample was used for LCMS analysis. The Bruker mass spectrometer was tuned for standard mass range analysis and data were continually acquired in the range *m/z* 50-3000; each run was recalibrated using a post-run injection of a sodium formate solution. Electrospray positive mode ionization was used with a source drying gas of 10 L/min at 200°C, nebulizer at 30 psi, and capillary set at 4000 V with an endplate offset of 500 V. For the Agilent Q-TOF, the Ion Funnel electrospray positive mode source used drying gas of 14 L/min at 200°C with nebulizer at 35 psi, a sheath gas flow of 11 L/min at 350°C, capillary set at 3500 V, nozzle voltage of 1000 V, and Fragmentor set at 175 V. Collision energies were set at 15 and 30 eV and data were continually acquired in the range *m/z* 50-1700 using a reference lock mass. Alkaloid responses were identified by retention times and high-resolution mass spectra. Although elemental composition determinations with less than 5 ppm accuracy were possible, and some MS/MS data were acquired, there is no LCMS database for frog alkaloid compounds. The occurrence of numerous alkaloid isomers made tentative identifications by LCMS challenging. Comparisons with GCMS data from frog skin were required for tentative identification of alkaloids given that standards for frog alkaloids are not commercially available.

### Arthropod genomic DNA extraction

Ant genomic DNA was isolated using a prepGEM Insect kit (ZyGEM, Hamilton, New Zealand) for smaller fragments or a NucleoSpin Tissue kit (Macherey-Nagel, Bethlehem, PA, USA) for full ants. Exoskeletons were crushed individually in a microcentrifuge tube with a sterile plastic pestle before following the manufacturers’ instructions. Mite genomic DNA was isolated using a phenol-chloroform extraction based on protocols from (Doyle 1987) and (Navajas et al. 1998) with some modifications. To crush the exoskeleton, each mite was pulverized individually in a microcentrifuge tube filled with 200 μL of extraction buffer (2% CTAB, 1.4 M NaCl, 0.2% 2-β mercaptoethanol, 20 mM EDTA, 100 mM TRIS-HCl pH 8) with a sterile plastic pestle before being incubated for 30 min at 60°C. Cold isopropanol was added to precipitate the DNA. After centrifugation, the DNA pellet was washed with 75% ethanol before resuspention in water. Purified genomic DNA was stored held at -20°C until PCR.

### Molecular identification of stomach contents

To identify mites and ants collected from the stomachs of *Oophaga sylvatica*, we used PCR to amplify the cytochrome oxidase 1 (CO1) region of genomic DNA, which is often used for arthropod taxonomic barcoding (Meusnier et al. 2008; Smith et al. 2005; Young et al. 2012). For all reactions, we used 2 μl of each primer (10μM) and 25 μL of 2X Phusion High-Fidelity PCR Master Mix with GC Buffer (New England Biolabs, Ipswich, MA) in a total reaction volume of 50 μL. Samples were amplified with either the primers LCO-1490 (5’-GGTCAACAAATCATAAAGATATTGG) and HCO-2198 (5’-TAAACTTCAGGGTGACCAAAAATCA) from (Folmer et al. 1994) or the primers CI-J-1632 (5’-TGATCAAATTTATAAT) and CI-N-2191 (GGTAAAATTAAAATATAAACTTC) from (Kambhampati and Smith 1995). A touchdown PCR program was used as follows: one round of 95°C for 5 min; 5 rounds of 95°C for 40 sec, 45°C for 40 sec with -1 °C per cycle, 72°C for 1 min; 40 rounds of 95°C for 40 sec, 40°C for 40 sec, 72°C for 1 min; one round of 72°C for 5 min. Successful reactions with one band of the expected size were purified with the E.Z.N.A. Cycle Pure Kit (Omega Bio-Tek, Norcross, GA) and Sanger sequenced by GeneWiz Inc. (Cambridge, MA).

We were able to PCR amplify CO1 in 45 out of 137 ant samples and 9 out of 20 mite samples. Nucleotide BLAST of the NCBI Genbank nr database was used to identify the resulting CO1 sequences (Online Resource Tables 1 and 2). For ants and mites, we assigned a family or genus level taxonomic identity based on the results of the BLAST search, where we considered greater than 96% sequence similarity sufficient to assign species or genera. For less than 95% similarity, we assigned specimens to a family based on BLAST similarity. For some ant specimens, we were able to identify the species based on the greater wealth of DNA barcoding information available for ants as compared to mites. For ants and mites separately, we used the software MEGA 6.0 (Tamura et al. 2013) to align using ClustalW the CO1 sequences from this study with other closely related species retrieved from GenBank. A nearest-neighbor joining tree was then constructed with a bootstrap of 5000 replications. All arthropod CO1 sequences have submitted to GenBank (Accession numbers: ants KU128453-97; mites KT947980-8).

**TABLE 1.**
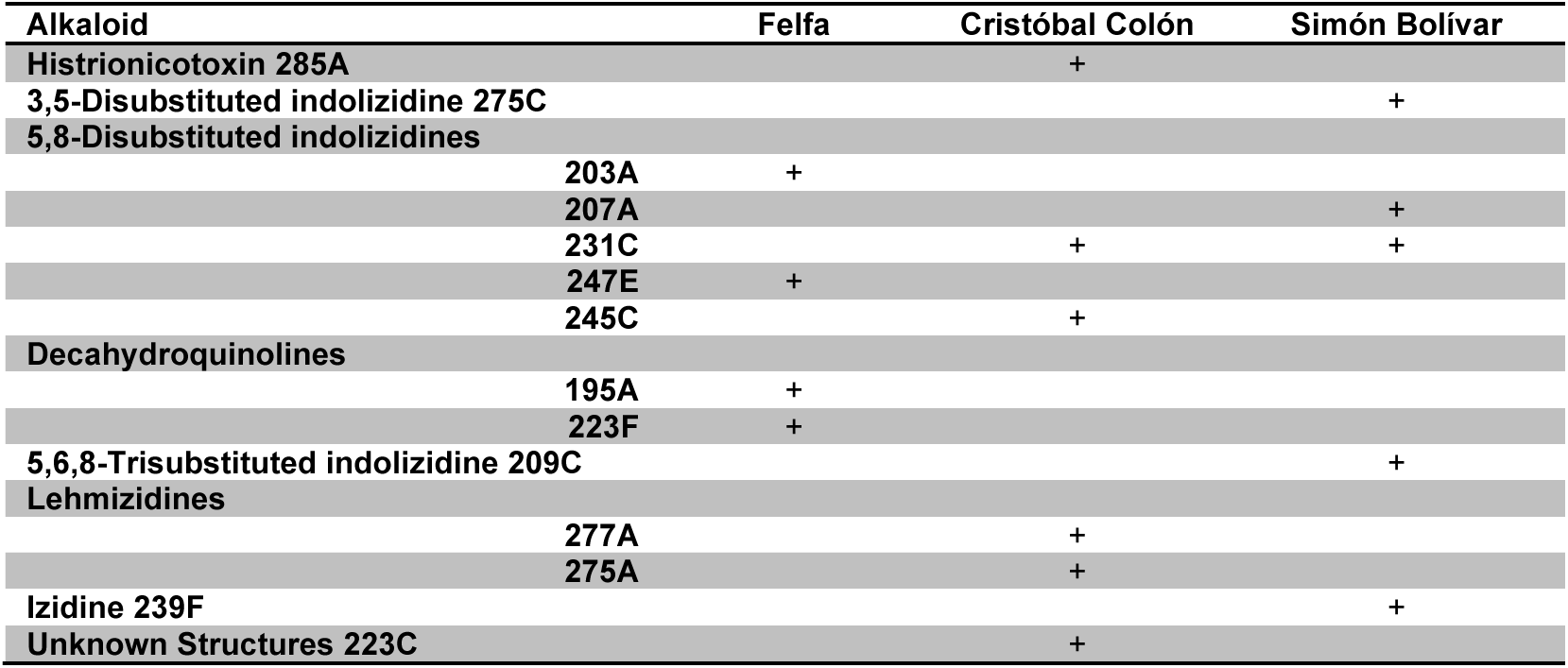
MAJOR ALKALOIDS IDENTIFIED IN *Oophaga sylvatica*

**TABLE 2.** BROAD DIET CHARACTERIZATION OF FELFA, CRISTÓBAL COLÓN, AND SIMÓN BOLÍVAR POPULATIONS

| Population | N | No. prey per frog<br>(mean (min & max)) | % ants<br>(num) | % ants<br>(vol) | % mites<br>(num) | % mites<br>(vol) | % beetles<br>(num) | % beetles<br>(vol) |
| --- | --- | --- | --- | --- | --- | --- | --- | --- |
| Felfa | 10 | 12.6 (6-33) | 78.72 | 85.81 | 15.86 | 6.825 | 3.693 | 7.317 |
| Cristóbal Colón | 10 | 9.6 (4-31) | 54.37 | 70.15 | 35.10 | 19.69 | 8.545 | 6.392 |
| Simón Bolívar | 12 | 12.5 (1-40) | 66.45 | 52.55 | 16.81 | 22.96 | 14.26 | 18.08 |

Abbreviations: N, number of frogs per population; num, number of arthropods; vol, volume of arthropods.

### Data analysis and statistics

GCMS analysis of alkaloids across populations relied on the extensive dataset by John Daly (Daly et al. 2005). Electron ionization mass spectrometry produced spectra of frog skin alkaloids with base peaks that are specific diagnostic ions based on the Daly GCMS database (Daly et al. 2005). Seven diagnostic ions were selected from their occurrence in the most abundant alkaloids across several frog populations. These extracted ion chromatograms were overlaid and normalized to the same scale to create a visual overview of major alkaloid distribution in different populations. Individual samples from each population were also processed in the same manner to visually compare individual variation within each frog population (See Online Resource). GCMS alkaloid data were statistically analyzed using an ANOVA and principal component analysis in XCMS software maintained by the Scripps Center for Metabolomics (Tautenhahn et al. 2012). XCMS calculates relative abundance by integrating the area under each ion current response. An ANOVA was run for each feature and a q-value (or Bayesian posterior p-value) was calculated to account for multiple testing (Storey 2003). All raw mass spectrometry data is available at DataDryad (TBD).

We used LCMS for analysis of alkaloids common to frogs and arthropods. For each frog skin, ant, and mite sample, five to seven of the most abundant alkaloids were selected for generation of an extracted ion chromatogram (EIC). Major ant or mite responses in the EIC were analyzed by molecular ion (M+H)+ product scans of both the ant or mite samples and the corresponding frog skin sample to compare both retention times and mass spectra. To confirm putative shared compounds with similar mass-to-charge ratios (m/z), 15 eV collision energy product ion scan total ion chromatograms (TIC) were compared to confirm the presence of a coeluting compound in both the frog skin and the arthropod sample of interest (ant or mite). Finally, we compared of 15eV product ion mass spectra from each putative compound shared by the frog and corresponding arthropod sample to confirm the spectra closely match, suggesting the compound of interest is identical in both frogs and arthropods.

To compare stomach contents across *O. sylvatica* populations, we quantified the number and volume of ants, mites, beetles, and other arthropods. Each diet variable was normally distributed, an ANOVA was used to determine population differences in different diet categories. A Tukey’s honest significance difference (HSD) test was used post-hoc to determine between population differences. A Benjimini Hochberg false discovery rate correction was applied to correct for multiple hypothesis testing.

## RESULTS

### Little Devil Frog Populations Vary in Alkaloid Profiles

We examined the alkaloid profiles from skin extracts of three *Oophaga sylvatica* populations located in northwestern Ecuador and compared them to a group of control (captive) frogs. Using GCMS, we found that each population had a unique profile of alkaloids (Figure 1a,b). Control frogs showed only three major peaks including D3-nicotine (spiked internal standard) and background from fatty acids. Principal component analysis shows distinct clustering between groups in principal component 1, which accounts for 21% of the alkaloid variation. The Cristóbal Colón population is centered between Felfa and Simón Bolívar (Figure 1c), which reflect their geographic distribution. Overall populations show robust between-population differences and little within-population variation (Online Resource Figures 1-3) with the exception of frogs collected from Cristóbal Colón. The most common compounds identified were 3,5-disubstituted indolizidines, 5,8-disubstituted indolizidines, decahydroquinolines, and lehmizidines (Table 1). Major alkaloids (the most abundant 5-6 alkaloids in each population) were mostly unique to a specific population, with overlap only between Cristóbal Colón and Simón Bolívar with the 5,8-disubstituted indolizidine 231C.

**Figure 1.**
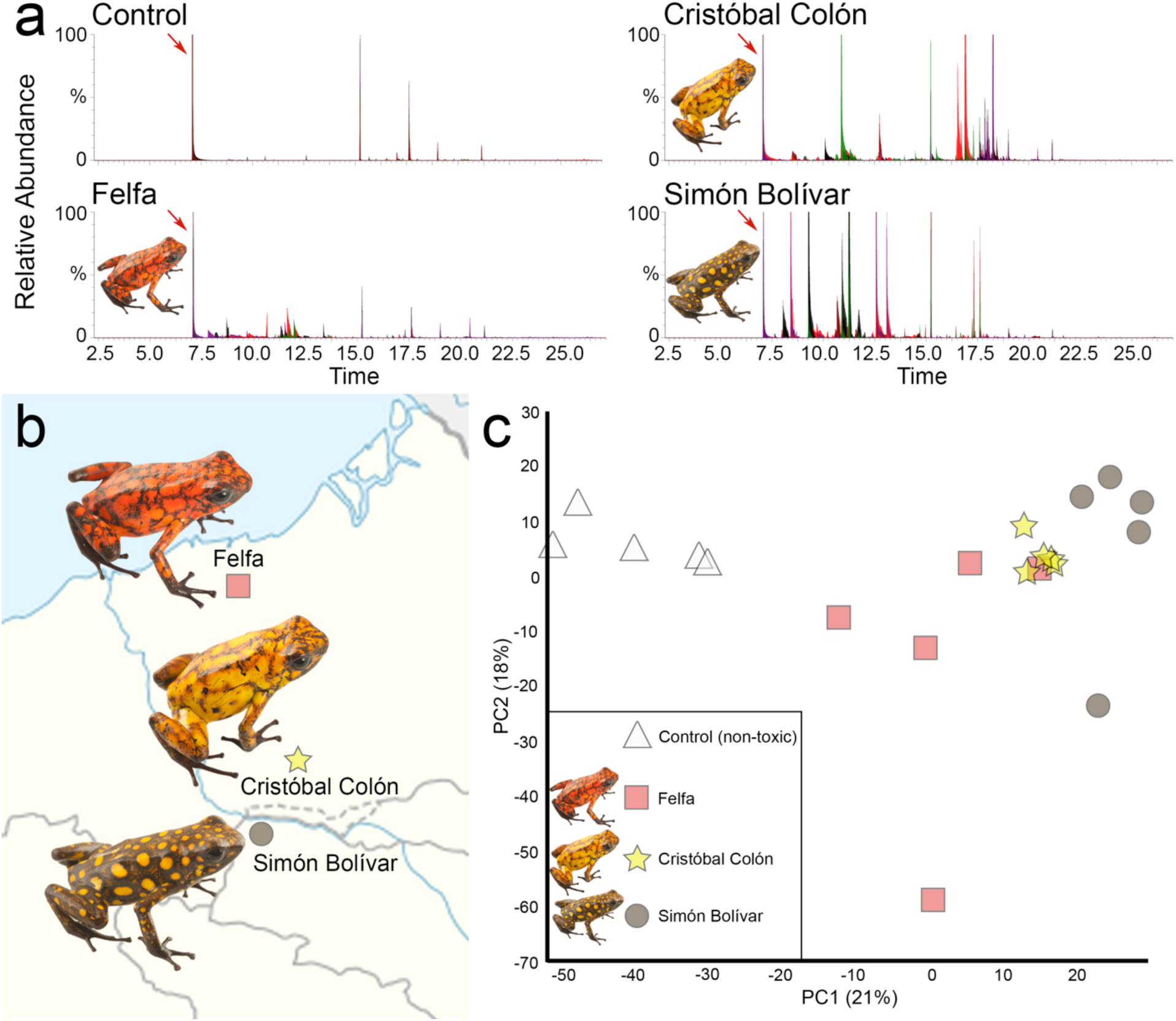
Little Devil frog *(Oophaga sylvatica)* population variation in chemical defenses. (a) Alkaloid toxin profiles assayed by gas chromatography mass spectrometry (GCMS) show population variation in chemical defenses where each peak is a chemical and peak amplitude represents abundance (N=5 per group). Overlaid colors represent different individuals (individual plots can be seen in supplementary materials). Note control (captive) frogs are rendered non-toxic with a diet of crickets and fruit flies; major peaks are skin fatty acids (not alkaloids) and benzocaine (used for euthanasia). Red arrows indicate spiked d3-nicotine as an internal reference. (b) Geographic distribution of the three *O. sylvatica* frog populations in northwestern Ecuador. (c) Principal component analysis of GCMS profiles reveals clustering of populations along a latitudinal gradient.

**Figure 2.**
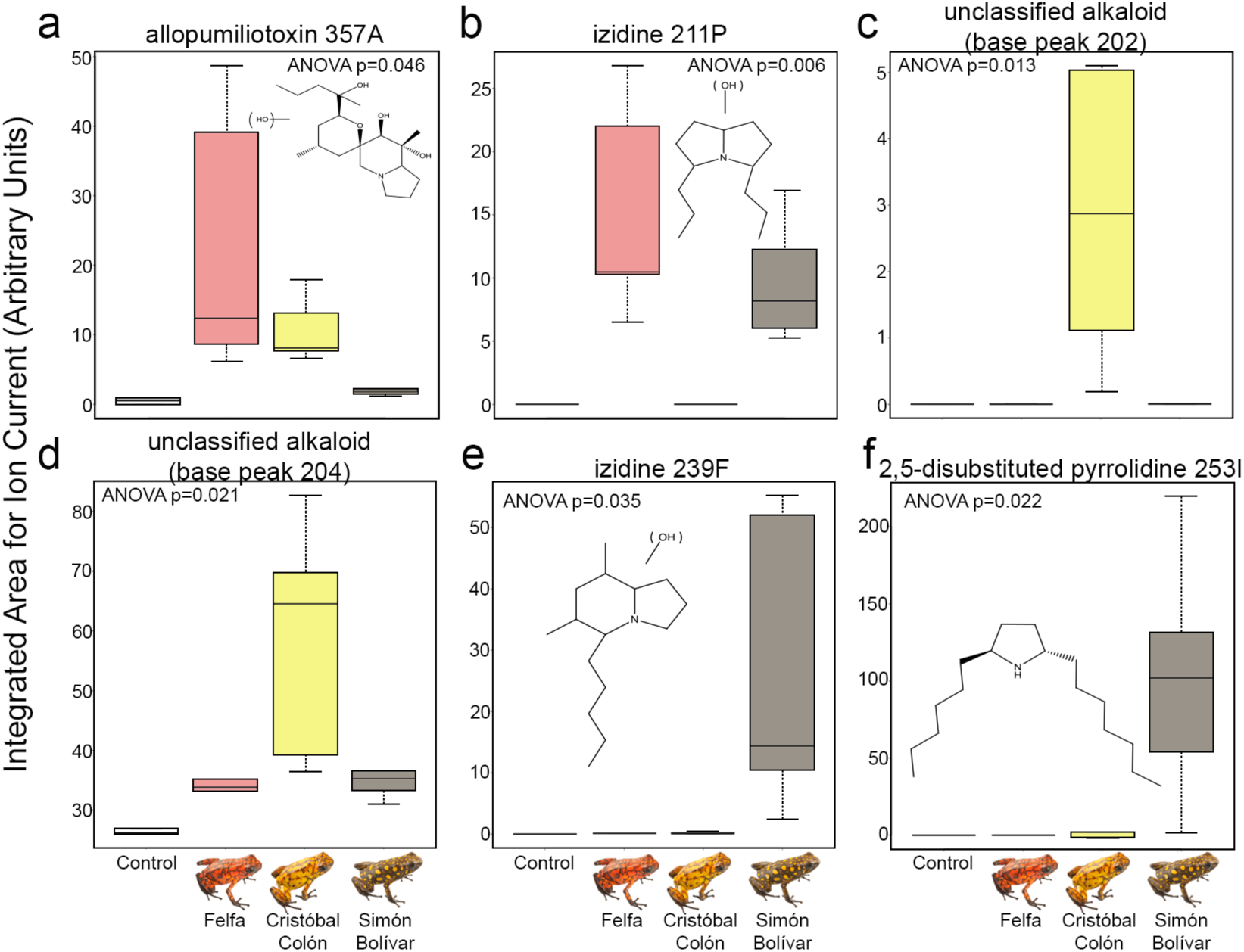
Each Little Devil frog *(Oophaga sylvatica)* population has a unique profile of defensive chemicals. Boxplots show relative abundance of various alkaloids between control (captive frogs; white) and wild populations of *O. sylvatica*, including Felfa (red), Cristóbal Colón (yellow), and Simón Bolívar (brown). GCMS extracted ion mass spectrum used for identification is shown in Supplementary Figure 4. Alkaloid structures are shown in each figure where known.

**Figure 3.**
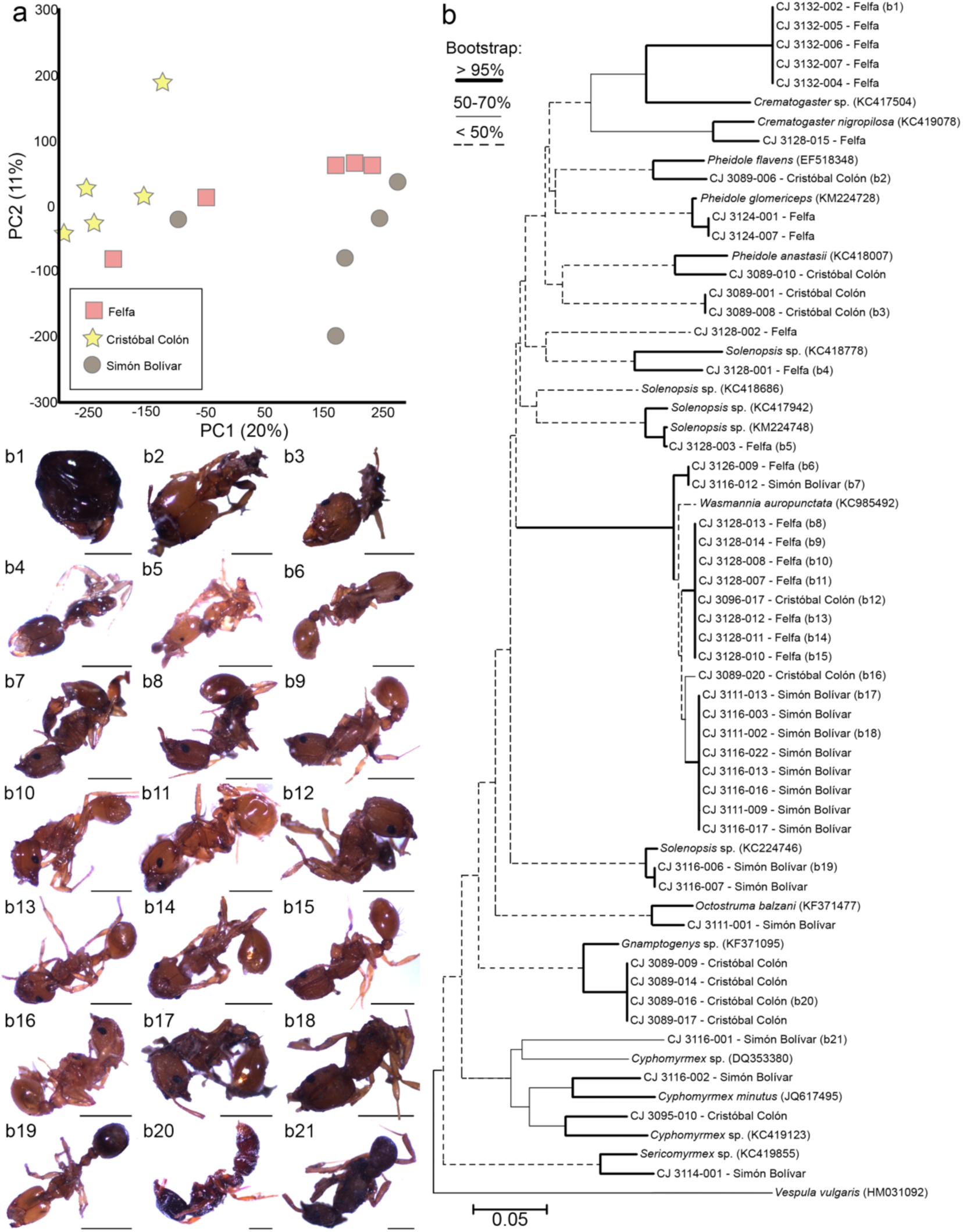
Identification and metabolomics of ants found in the stomachs of Little Devil frogs *(Oophaga sylvatica)*. (a) Principal component analysis clustering of ant liquid chromatography mass spectrometry data colored by frog population. (b) A phylogenetic tree with nearest neighbor clustering shows relationships in cytochrome oxidase 1 sequence between ants isolated from *O. sylvatica* stomachs compared to other known ants. Ants are numbered by the four-digit frog specimen identifier followed by a three-digit number assigned to the arthropods in the order they were isolated from a single stomach. The frog population each ant was isolated from is listed beside each identifier. Genbank IDs of known ants are shown after the species name. The common wasp *(Vespula vulgaris)* was used as an outgroup. (b1–21) Photos of some specimen are shown, with varying degrees of being intact or partially digested. Scale bar is 5 mm.

We examined in more detail the identity of alkaloids that differed in abundance between frog populations but were absent in control frogs fed crickets in captivity (Figure 2, Online Resource Figure 4). Two alkaloids were highly abundant in the Felfa population, and were shared with one of the other populations surveyed. Allopumiliotoxin 357A is present in both Felfa and Cristóbal Colón (ANOVA, *F*_3,20_ = 5.278, P = 0.046) (Figure 2a) and izidine 211P is present in Felfa and Simón Bolívar populations (ANOVA, *F*_3,20_ = 11.439, P = 0.006) (Figure 2b). Two unclassified alkaloids were more abundant in the Cristóbal Colón population. One unclassified alkaloid (Figure 2c) has a base peak of 202 (Online Resource Figure 4c), characteristic of some unclassified alkaloids (ANOVA, *F*_3,20_ = 8.192, P = 0.013). The second unclassified alkaloid (Figure 2d) has a base peak of 204 (Online Resource Figure 4d), a characteristic of many unclassified alkaloids (ANOVA, *F*_3,20_ = 6.809, P = 0.021). Two alkaloids were abundant in the Simón Bolívar population including izidine 239F (ANOVA, *F*_3,20_ = 5.798, P =0.035) and trans-2-hexyl-5-heptyl-pyrrolidine (253I; ANOVA, *F*_3,20_ = 6.615, P = 0.022) (Figure 2e-f; Online Resource Fig 4e-f). Overall, our results show a unique profile of alkaloid toxins within each population, with little overlap between populations. This observation was replicated with LCMS data (Online Resource Figure 5).

**Figure 4.**
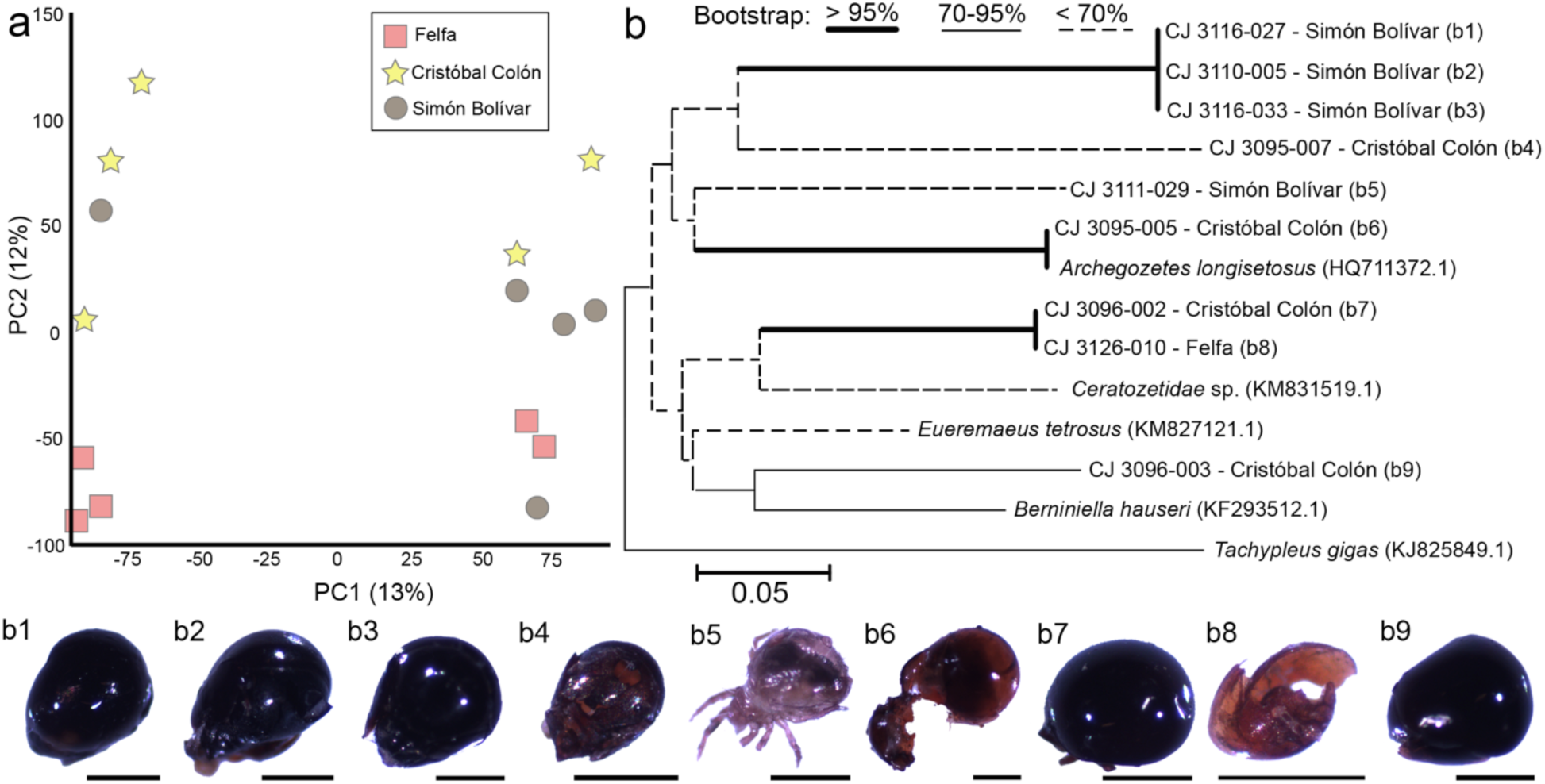
Identification of mites found in the stomachs of Little Devil frogs *(Oophaga sylvatica)*. (a) Principal component analysis clustering of mite liquid chromatography mass spectrometry data colored by frog population. (b) A phylogenetic tree with nearest neighbor clustering shows relationships in cytochrome oxidase 1 sequence between mites isolated from *O. sylvatica* stomachs compared to other mites. Isolated mites are numbered by the four-digit frog identifier followed by a three-digit number assigned to the arthropods in the order they were isolated from a single frog stomach. The frog population each mite was isolated from is listed beside each identifier. Genbank IDs of known mites are shown after the species name. A horseshoe crab *(Tachypleus gigas)* was used as an outgroup. (b1–9) Photos of each specimen are shown, with varying degrees of being intact or partially digested. Scale bar is 5 mm.

**Figure 5.**
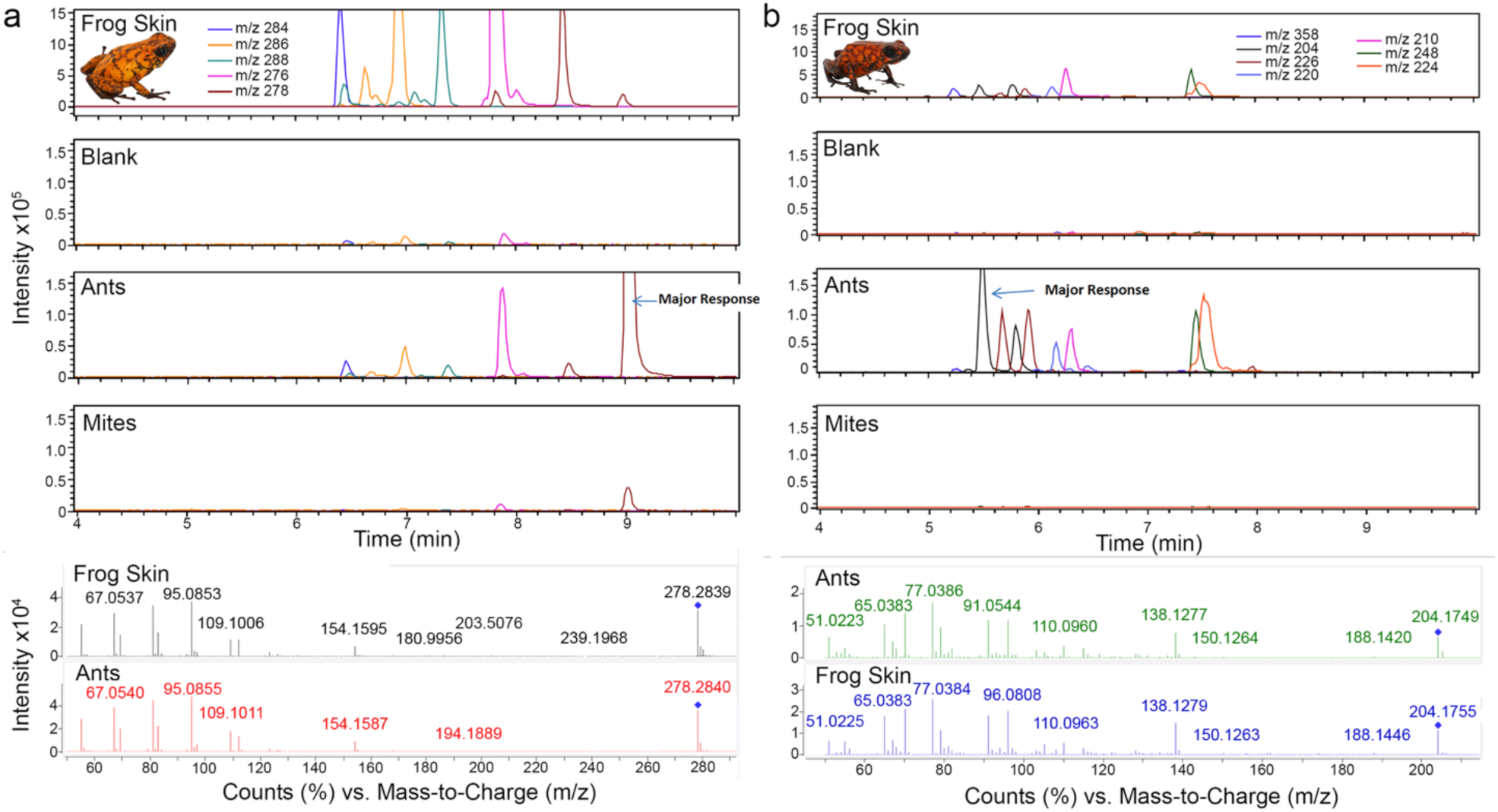
Identification of ants as a dietary source of Lehmizidine 277A and 5,8-disubstituted indolizidine 203A in the Little Devil frog *(Oophaga sylvatica)*. Liquid chromatography mass spectrometry (LCMS) was used to analyze alkaloids from *O. sylvatica* skin samples and the ants and mites found in those same frogs' stomachs. Top: Abundant alkaloids in the frogs were selected for extracted ion chromatogram analysis. Each compound is represented as a different color and mass-to-charge (m/z) ratios are listed at the top of each panel. In both comparisons from two different frogs, a major alkaloid response was detected in the ant sample that was similar to a response found in the frog skin sample. Bottom: Tandem mass spectrometry fragmentation patterns via 15 eV product ion mass spectra shows a close match between the compounds found in the frogs and the ants. (a) A similar response of m/z 278 was found in a Cristóbal Colón frog skin sample and the corresponding ant sample. Comparison with the GCMS data from this frog suggests the identity of this shared alkaloid is lehmizidine 277A. (b) A similar response of m/z 204 was found in a Felfa frog skin sample and the corresponding ant sample. Comparison with the GCMS data from this frog suggests the identity of this shared alkaloid is 5,8-disubstituted indolizidine 203A.

#### Dietary Differences Between Little Devil Frog Populations

To gain a better understanding of how diet contributes to alkaloid profiles in these Little Devil frog populations, we determined the quantity and identity of arthropods collected from the stomach contents using both morphometric and molecular methods. We first grouped them into broad categories of ants, mites, beetles, and "other” (Table 2) and calculated the number and volume of each arthropod type. With a subset of ants and mites from each population (N=5 frogs per population), we used an untargeted metabolomics approach with LCMS to characterize the chemical signature of consumed arthropods across frog populations. For the remaining ants and mites, we sequenced the CO1 gene to characterize species diversity.

#### Species and chemical diversity in consumed ants

When characterizing broad patterns in ant consumption across frog populations (Table 2), we found that the percentage of ants by volume varied between populations (ANOVA, *F*_2,32_ = 4.480, P = 0.020), but the percentage of ants by number did not. Felfa had a higher percentage of ants by volume compared to Simón Bolívar (Tukey’s HSD, P = 0.015) but not compared to Cristóbal Colón (Tukey’s HSD, P = 0.382). Cristóbal Colón and Simón Bolívar did not differ in the percentage of ants by volume.

To determine if consumed ants (pooled by individual frog) vary in their chemical profiles across Little Devil frog populations, we performed a principal component analysis of ant LCMS data (Figure 3a). We found that prey ant chemical profiles also cluster by the frog population from which they were isolated, suggesting the three *O. sylvatica* frog populations have access to different ant prey with distinct chemical profiles.

To better understand how ant prey may differ in these geographically distinct frog populations, we obtained CO1 sequence for 45 out of 137 ant specimens isolated from frog stomachs. Consumed ants belong to seven genera of the Myrmicinae sub-family (tribe Attini [fungus-growing ants], genera: *Cyphomyrmex, Octostruma, Pheidole, Sericomyrmex, Wasmannia;* tribe Crematogastrini, genus: *Crematogaster* [acrobat ants]; tribe Solenopsidini, genus: *Solenopsis* [stinging ants]) and from one genus of the Formicidae Ectatomminae subfamily (tribe Ectatommini, genus: *Gnamptogenys* [predatory ants]) (Figure 3b, Online Resource Table 1). All the samples have more than 80% sequence similarity with their top BLASTn hit matches and 22 have more than 96% similarity, sometimes allowing resolution at the species level. However, some specimens have close BLASTn matches (80-90%) for different ant genera (usually undescribed species), which do not allow confident assignment of some specimens. Many specimens only had a match of up to 85% homology in CO1 sequence to records in the GenBank database, suggesting these may be undescribed ant species.

Based on CO1 sequence similarity between ant specimens, we estimate 20 different species of ants were recovered. Two genera of *Myrmicinae Attini* ants were recovered from frog stomach contents across all three *O. sylvatica* populations, including *Solenopsis* and *Wasmannia*. A unique species to the Felfa group is a *Myrmicinae Crematogastrini Crematogaster* species, which was not observed in the two other frog populations. These *Crematogaster* ants were found in the same Felfa frog (CJ 3132), and the closest match in the GenBank database is only 86% similarity, suggesting these specimens represent an undescribed ant species. A different *Crematogaster* species was isolated in a separate Felfa frog (CJ 3138) and has 94% similarity in CO1 sequence with *Crematogaster nigropilosa*. Four ant specimens collected from both Felfa (CJ 3124) and Cristóbal Colón (CJ 3089) cluster with *Myrmicinae Attini Pheidole* genera, although there is not an exact match with any described ant CO1 sequences in the GenBank database. Finally, four specimens recovered from a Cristóbal Colón (CJ 3089) are the only *non-Myrmicinae* ants we recovered. These specimens have identical CO1 sequence with each other, and are 95% similar to the *Ectatomminae Gnamptogenys* genera.

#### Species and chemical diversity in consumed mites

The amounts of mites consumed between populations varied (Table 2) The percentage of mites in the stomach contents by number differed between populations (ANOVA, *F*_2,32_ = 4.232, P = 0.024) where the Cristóbal Colón population had a higher percentage of mites by number compared to both Felfa (Tukey’s HSD, P = 0.042) and Simón Bolívar (Tukey’s HSD P = 0.044). The percentage of mites by volume also varied between populations (ANOVA, *F*_2,25_ = 7.063, P = 0.004), where Simón Bolívar frogs had a higher percentage of mites by volume in the stomach contents compared to Felfa (Tukey’s HSD, P = 0.003).

To determine if the chemical profiles of consumed mites were distinct between frog populations, we performed a principal component analysis of the mite LCMS data (Figure 4a). We found clustering of mite samples according to frog population in principal component 2 (12% of variance) but not principal component 1 (13% of variance).

We hypothesized that diversity in consumed mite species was responsible for the clustering of chemical profiles observed in the principal component analysis. To test this hypothesis, we obtained CO1 sequence for 9 out of 20 mites individually isolated from frog stomachs. Based on the top BLASTn hit for each mite, all specimens are likely Oribatid (Order Oribatida) mites (Figure 4b, Online Resource Table 2). However, due to the low percent similarity in CO1 sequence of most mites to the mites in the GenBank nr database (78-84%), we were unable to assign mites to particular genera and many of these specimens likely represent undescribed species. An exception is one mite that has a 99% similarity to the mite *Archegozetes longisetosus*, and is likely the same species. Additionally three mites isolated from two Simón Bolívar frogs had identical morphology and CO1 sequence, suggesting they are the same undescribed species. Thus, we have identified 5 undescribed and 1 described species of mites from the diet of three *Oophaga sylvatica* populations. Unfortunately, we did not obtain enough sequence information from separate mite specimens to conclusively determine how species diversity drives chemical diversity observed in the mite LCMS data.

#### Other dietary arthropods

In addition to ants and mites, which together represent over 80% of the *Oophaga sylvatica* diet, we also evaluated the number and volume of beetles and "other” arthropods isolated from frog stomach contents (Table 2). However, the number of arthropods in these categories was so low that we did not evaluate statistical differences between frog populations. Nearly all of the frogs from Cristóbal Colón had eaten at least one beetle (8 out of 10 frogs) whereas only 2 Simón Bolívar and 3 Felfa frogs had eaten a beetle. Very few arthropods were isolated that did not group into the categories of ants, mites, or beetles (2 in Felfa, 5 in Simón Bolívar, and 4 in Cristóbal Colón). These specimens were usually flies, wasps, or stinkbugs, although we did not use molecular methods to identify these samples. We did not examine the chemical profiles of non-mite or non-ant arthropods using LCMS.

### Poison Frog Alkaloids in Ants and Mites

While the dietary specialization of poison frogs on ants and mites has been well documented, less is known about which alkaloid compounds are derived from specific prey categories. We compared three samples (frog skin, ants, and mites) for each of five individuals from each population using LCMS, which gives increased sensitivity compared to GCMS. By comparing all three samples within each individual, we were able to trace the dietary source of four alkaloids.

#### Alkaloids Detected in Ants

We compared the LCMS responses in frog skin and their corresponding prey samples in search of compounds that had similar mass to charge ratios in both the frog skin and the ants, but not the mites. We then further examined these compounds with tandem mass spectrometry to confirm the alkaloids found in frog skin and ants had similar fragmentation mass spectra patterns, which suggests the compound of interest is identical in both samples.

Using this approach we identified ants as the dietary source of two frog alkaloids (Figure 5; Online Resource Figure 6). In a Cristóbal Colón frog skin (CJ 3093) and the corresponding sample of consumed ants (Figure 5a), we observed a compound with a mass to charge ratio of 278 that also had identical fragmentation patterns in both samples, suggesting these compounds are identical. Comparing the LCMS data to the GCMS data from the same frog skin sample, we conclude that this ant-derived alkaloid is lehmizidine 277A. Similarly, we identified a response in a Felfa frog (CJ 3129) and consumed ants from that same frog (Figure 5b). This compound has a mass to charge ratio of 204, and the fragmentation patterns show the mass spectra closely match, suggesting identical compounds in both frog and ant samples. From comparison of GCMS responses with m/z 204 in this same frog skin sample, this alkaloid is likely 5,8-disubstituted indolizidine 203A.

**Figure 6.**
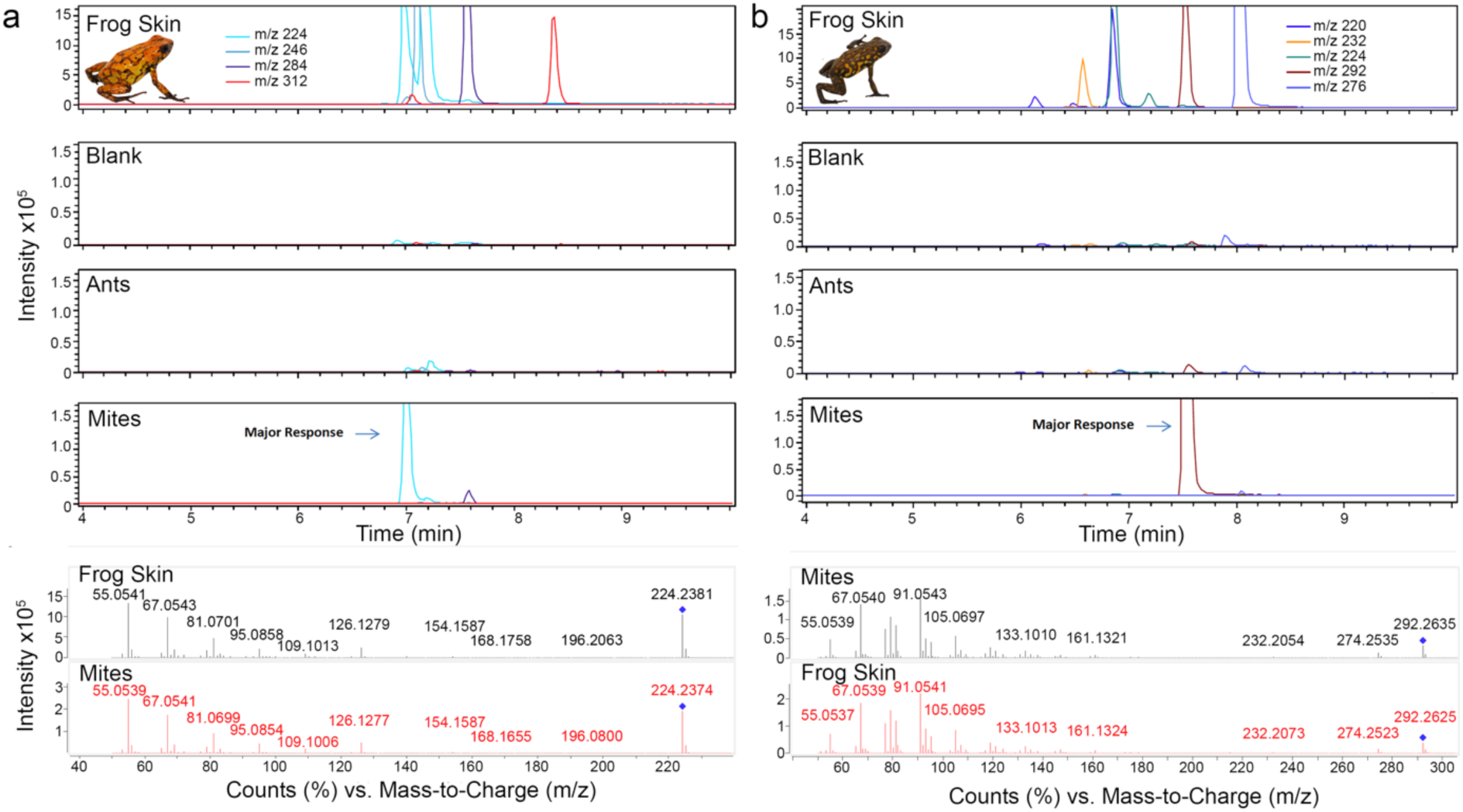
Identification of mites as dietary source of 3,5-disubstituted indolizidine 223AB and an unknown alkaloid in the Little Devil frog *(Oophaga sylvatica)*. Liquid chromatography mass spectrometry (LCMS) was used to analyze alkaloids from O. sylvatica frog skin samples and the ants and mites found in those same frogs' stomachs. Top: Abundant alkaloids in the frogs were selected for extracted ion chromatogram analysis. Each compound is represented as a different color and mass-to-charge (m/z) ratios are listed in the top of each frog panel. In both comparisons from two different frogs, a major alkaloid response was detected in the mite sample that was similar to a response found in the frog skin sample. Bottom: Tandem mass spectrometry fragmentation patterns via 15 eV product ion mass spectra shows a close match between the compounds found in the frogs and the mites. (a) A similar response of m/z 224 was found in a Cristóbal Colón frog skin sample and the corresponding mite sample. Comparison with the GCMS data from this frog suggests the identity of this shared alkaloid is 3,5-disubstituted indolizidine 223AB. (b) A similar response of m/z 292 was found in a Simón Bolívar frog skin sample and the corresponding mite sample. Identification of this alkaloid is not possible, as it was not observed in GCMS data for this frog.

#### Alkaloids Detected in Mites

We took a similar approach to detect common LCMS responses in frog skin and their corresponding mite samples, where compounds should have similar mass to charge ratios in both the frog skin and the mites, but not the ants. Further examination of these compounds with tandem mass spectrometry confirmed the alkaloids found in frog skin and mites are identical.

Using this comparative LCMS approach, we identified mites as the dietary source of two poison frogs alkaloids (Figure 6, Online Resource Figure 6). In a Cristóbal Colón frog (CJ 3091), a similar LCMS response with a mass to charge ratio of 224 was observed in both the frog skin and the pooled mite sample (Figure 6a). Comparison of the tandem mass spectrometry fragmentation pattern shows the spectra closely match, suggesting the compounds detected in these samples are identical. From comparison of the GCMS responses with a mass to charge ratio of 224 in this same frog skin sample, this compound is likely 3,5-disubstituted indolizidine 223AB. In a separate frog-mite comparison, a similar LCMS response with a mass to charge ratio of 292 was observed in a Simón Bolívar frog (CJ 3112) and the pooled mites from the frog’s stomach (Figure 6b). Fragmentation mass spectra of the compounds suggest this compound is identical. From the GCMS analysis of the frog skin sample, no corresponding alkaloid with a similar mass to charge ratio was detected, likely from the lack of sensitivity of this method compared to LCMS.

## DISCUSSION

We have shown that *Oophaga sylvatica* populations vary in defensive chemical profiles in a manner that reflects prey arthropod chemical diversity. We used mass spectrometry to identify the arthropod source of four alkaloid toxins by comparing frog skin chemistry to arthropods isolated from the stomach contents of the same frogs. Overall our results highlight the diversity of chemical defenses found in *O. sylvatica* and Ecuadorean ants and mites, adding to the growing literature of the trophic relationships of poison frog toxins and their arthropod sources.

### Little Devil frog defensive chemicals

We used GCMS to determine the repertoire of defensive chemicals across three *Oophaga sylvatica* populations. We observed in a principal component analysis of frog alkaloid data that populations clustered along their latitudinal gradient. Research on poison frog population differences in alkaloid toxins have recently focused on the Strawberry poison frog (*Oophaga pumilio)* in Costa Rica and Panama (Saporito et al. 2006; Saporito et al. 2007a; Saporito et al. 2010; Saporito et al. 2012), and to a lesser extent the Imitator frog *(Ranitomeya imitator)* in Peru (Stuckert et al. 2014). These studies have also shown greater similarity in alkaloid profiles with geographic proximity (Saporito et al. 2007a). Variation in alkaloid profiles among populations has been described for some *O. sylvatica* populations (under the name *Dendrobates histrionicus)*, as well as a number of other *Oophaga* species, including *O. histrionica* and *O. lehmanni* (Myers and Daly 1976). Dietary environment seems to be a significant component of toxin profiles within *Oophaga*. For example, sympatric populations of *O. granulifera* and *O. pumilio* have more similar alkaloid profiles than the geographically distant populations of *O. granulifera* (Myers et al. 1995). However, this is not always the case as co-mimetic (sympatric) species of *Ranitomeya* differ in their alkaloid profiles (Stuckert et al. 2014), which may suggest variation in micro-habitats, contrasting prey preferences, or genetic differences that impact the efficiency and specificity of alkaloid sequestration. Clearly more comparative work across broader range of species and habitats are required to fully understand how these ecological and genetics factors impact frog chemical defenses.

The most abundant classes of alkaloids in *Oophaga sylvatica* were 3,5- or 5,8-disubstituted indolizidines, although the precise alkaloid in these classes differed between the three populations (Table 1). This is in contrast to toxin profiles for this species collected from Ecuador by Myers and Daly (Daly et al. 1978) who reported only histrionicotoxins as the major alkaloids in several populations, which we only detected in Cristóbal Colón frogs. This discrepancy could be due to sampling location as *O. sylvatica* in the (Myers and Daly 1976) study were collected in southernmost part of their range. Other possibility is could be changes in dietary arthropod composition over the past 40 years, which may have shifted the most abundant alkaloids in this species from histrionicotoxins to disubstituted indolizidines. The unique major alkaloids in Cristóbal Colón compared to Felfa and Simón Bolívar include histrionicotoxin 285A and lehmizidines 277A and 275A. Only Felfa frogs had decahydroquinolines (DHQ 195A and 223F) as a major alkaloid group. Many of the alkaloids we have identified in *O. sylvatica* have also been detected in *O. pumilio* (Saporito et al. 2007a), with the exception of 5,8-I 245C and Izidine 239F, and some *Mantellas* (Garraffo et al. 1993).

Individual variation in alkaloid profiles was limited in the Simón Bolívar and Felfa populations, but was much greater in the Cristóbal Colón population where almost all frogs had strikingly different alkaloid profiles. The general diet of Cristóbal Colón frogs did not drastically differ and without genetically profiling their stomach contents (which were instead used for LCMS quantification of alkaloids), we are unable to determine the species diversity of their diets, which we predict from the alkaloids profiles to be very different. However, it should be noted that quantification of stomach contents represents a snapshot in time and may not be representative of the full dietary repertoire that contributed to skin alkaloid profiles accumulated over weeks or months. Additionally, frogs that had high alkaloid peaks around 16-19 minutes by GCMS were all male, whereas the other two frogs with low alkaloid levels around this time point were female. It could be that there are sex differences in toxin sequestration within the population, as has been documented in some *O. pumilio* populations (Saporito et al. 2010). However, this is unlikely as both males and females were included in the Simón Bolívar and Felfa populations, which did not show a high degree of individual variation within the population. There are two more plausible explanations for this individual variation. First, frog age may be a contributing factor to alkaloid diversity as was recently documented in the Brazilian red-belly toad *(Melanophryniscus moreirae)* (Jeckel et al. 2015). Unfortunately, we were unable to determine the age of the *O. sylvatica* frogs, although this is likely an important variable to consider in future studies. Secondly, frogs may have been collected in different microhabitats and this difference in diet availability may influence alkaloid profiles. This has been recently observed in *Mantellas* where frogs from disturbed habitats have a higher diversity in alkaloids than frogs from undisturbed habitats (Andriamaharavo et al. 2010). Further sampling and broader analyses of potential contributing factors within the Cristóbal Colón population would be required to fully understand these individual differences in alkaloid profiles.

### Arthropod diversity in the diet of poison frogs

The poison frog dietary specialization on ants and mites is well established (Caldwell 1996; Darst et al. 2005). As poison frogs sequester their toxins from their diet, it has been proposed that the diversity of poison frog toxins reflects the diversity of arthropods in the tropics (Saporito et al. 2009). We found that the stomach contents of the three *Oophaga sylvatica* populations contained over 80% ants and mites by number. Ants in particular composed at least 50% of the diet in these populations. Our results are similar to those of *O. histrionica* (Osorio et al. 2015), a closely related species in Colombia, whose diet is mainly composed of Formicidae (ants), Acari (mites), and Coleoptera (beetles). These three arthropod families account for over 97% of the diet (by number) of the three *O. sylvatica* populations we investigated. We found that Felfa frogs tended to have more ants in the stomach contents, and ants have been proposed to contain mostly unbranched alkaloids (Saporito et al. 2012). We also identified DHQs 195A and 223F as major alkaloids in Felfa, which have an ant origin (Jones et al. 1999). On the other hand, both Cristóbal Colón and Simón Bolívar populations had higher proportion of mites in their stomach contents (either by number or volume, respectively), which have been proposed as the main source of branched-chain alkaloids (Saporito et al. 2007b; Saporito et al. 2011).

*Ants*. Ant genera diversity and chemical variation reflects the heterogeneity of *Oophaga sylvatica* toxin profiles across populations. We found that ant metabolomics data clustering by frog population in a principal component analysis, suggesting geographic differences in toxin availability within frog prey items. We identified ant genera in *O. sylvatica* stomachs that are often consumed by other poison frogs such as *Pheidole, Crematogaster, Cyphomyrmex, Octostruma, Gnamptogenys, Solenopsis and Wasmannia* (Gómez-Hoyos et al. 2014; Mebs et al. 2014; Osorio et al. 2015). Most of the consumed ants were in the Subfamily *Myrmicinae* Tribe *Attini*, the fungus-cultivating ants. Specimens in the genus *Solenopsis* were present in all frog populations, but likely represent different species or subspecies given their diversity in CO1 sequence and morphology. Recently the genus *Solenopsis* was identified as a source for histrionicotoxin (Jones et al. 2012), and we detected the presence histrionicotoxin in Cristóbal Colón frogs that also ate *Solenopsis* ants. However it should be noted that although *Solenopsis* species and histrionicotoxins were observed in Cristóbal Colón, we did not detect histrionicotoxins in Felfa or Simón Bolívar frogs, which also ate *Solenopsis* ants. This discrepancy is a caution against assuming every *Solenopsis* ant species is chemically similar or that every ingested arthropod found in a poison frog stomach contributes to the alkaloid repertoire of the frog. Finally, *Pheidole* ants are also known to be predators of Oribatid mites (Wilson 2005), which can carry alkaloid toxins also identified in frogs. This raises the possibility of toxin transfer from mites through ants to frogs, as well as from mites to frogs directly (Saporito et al. 2011).

*Mites*. We found the genera and chemical diversity of mites partially reflects the variation in toxin profiles across *Oophaga sylvatica* populations. We sequenced the CO1 region from 9 mite specimens representing 6 undescribed species. We initially isolated 20 mites from frog stomachs, but the low success rate of CO1 amplification is likely due the small amount of genomic DNA we were able to extract from these tiny arthropods. The closest match in the GenBank nr database for all samples are Oribatid mites, which are a source of many poison frog alkaloids (Saporito et al. 2007b). Interestingly, Saporito et al. (Saporito et al. 2011; Saporito et al. 2015) has shown Oribatid mites carry many alkaloids found on poison frogs as well as many other non-frog alkaloids. A recent study has shown tropical mites carry more alkaloids than temperate mites (Saporito et al. 2015), although it is not clear if poison frog alkaloid variation reflects diversity in Oribatid mite species across a smaller longitudinal gradient. Our principal component analysis of mite metabolomics data shows only partial clustering by frog populations, but more chemical profiling of mites is needed to determine how they contribute to geographic variation in poison frog toxin profiles. Interestingly, Oribatid mites seem to vary in alkaloid profiles based on developmental stage, and (Saporito et al. 2011) suggest this may indicate mites synthesize these alkaloids themselves. However, only plants, bacteria, and fungi have been demonstrated to have the enzymatic pathways necessary to synthesize alkaloid neurotoxins (Ashihara et al. 2008; Osswald et al. 2007; Waller 2012; Wiese et al. 2010), and so it remains to be determined if mites themselves synthesize alkaloids or if they obtain these small molecules from plants or microbes. Regardless of the alkaloid origin in mites, they are clearly a rich biochemical resource for poison frog chemical defense and should be studied in this context more systematically.

### Tracing the source of poison frog toxins

Since the dietary hypothesis of poison frog toxicity was proposed in the 1990s (Daly et al. 1994a; Daly et al. 1994b), there has been a strong focus on identifying the arthropods harboring these alkaloid chemicals. Many of the alkaloid classes have been identified in either ants or mites (or both) (Santos et al. 2015; Saporito et al. 2012; Saporito et al. 2009) found sympatry with poison frogs. For example, histrionicotoxins and pumiliotoxins have been identified in ants (Jones et al. 2012; Saporito et al. 2004). However, it has been historically difficult to identify poison frog alkaloids in arthropods given their small size and low alkaloid content per specimen as well as difficulty in obtaining species that poison frogs have been known to ingest from the leaf litter, especially extremely small mites.

To identify the arthropod source of *Oophaga sylvatica* toxins, we took a simple approach of pooling separately the ants and mites found in the frogs’ stomachs. This allowed us to examine the chemistry of arthropods the frogs ingested rather than surveying the general surrounding leaf litter that may contain many arthropods the frogs do not eat. Moreover, the high sensitivity of LCMS allowed us to detect small quantities of alkaloids that may be undetectable by GCMS. However, because the alkaloid data library by Daly and colleagues is based on GCMS results (Daly et al. 2005), examining frog skin alkaloids must be done with both GCMS (for alkaloid identification) and LCMS (for comparison with arthropod data). This dual approach allowed us to identify the arthropod source of three alkaloids (277A, 203A, and 223AB), but for one alkaloid we were not able to tentatively assign an identity given it was not at detectable levels using GCMS. It is important to note that our experimental design for LCMS limits our ability to identify the exact species of ant or mite from which alkaloid is found. Although we photographed the arthropods prior to alkaloid extraction, we cannot reliably identify the species within the pooled sample without genetic testing. New methods involving DNA barcoding of fecal samples to characterize diet (Kartzinel and Pringle 2015) may be a useful tool in the future to genetically identify small arthropod prey without sacrificing frogs for stomach content analysis. Finally, it is important to note that the LCMS profiling of stomach contents represents a snapshot in time. Many frog alkaloids were not detected in the arthropod samples given they do not represent the full repertoire of dietary diversity in these frogs. The pharmacokinetics of toxin sequestration in poison frogs is unknown, and some frog toxins could have been acquired weeks to months prior to our sampling. A combination of non-invasive and repeated sampling of prey arthropods and frog skin toxins would be a good step forward to resolving some of the pitfalls of standard methods in this field.

To our knowledge, we have provided the first evidence of the dietary source of three alkaloid toxins. We found lehmizidine 277A in frogs of the Cristóbal Colón population and traced the dietary source of this toxin to ants. Ants have been the proposed source of lehmizidines for poison frogs since monosubstituted lehmizidines occur in a myrmicine ant (Jones et al. 2007), but we provide here the first evidence for this trophic relationship. We were also able to identify ants as the source of 5,8-disubstituted indolizidine 203A in frogs from Felfa. Other alkaloids in this class have been identified in both ants and mites (Daly et al. 2002; Saporito et al. 2007b). Finally, we were able to trace the source of 3,5-disubstituted indolizidine 223AB in Cristóbal Colón frogs to mites. Other alkaloids in this class have also been attributed to both ants and mites (Jones et al. 1999; Saporito et al. 2007b). Finally, although we were able to identify an alkaloid (m/z 292) in both frog skin from Simón Bolívar and mites found in that same frogs’ stomach, we were unable to tentatively identify the alkaloid given the lack of LCMS library data for these chemicals. It is the partnership between biologists and chemists (largely John Daly, (Savitzky and Saporito 2012)) that has driven the field of chemical ecology in poison frogs forward. With emerging new technologies, such interdisciplinary partnerships are still necessary for progress.

### Summary

We have described here the alkaloid profiles of three populations of *Oophaga sylvatica*, the arthropods that compose their distinctly different diets, and have identified the dietary source of four poison frog alkaloids. Moreover, our results highlight how arthropod species and chemical diversity drives variation in poison frog chemical defenses. Future work will focus on identifying the arthropod species that harbor these alkaloids, tracing the source of these toxins to the plants or microbes that synthesize them and profiling additional *O. sylvatica* populations.

## Acknowledgements

We are grateful to Jon Clardy, Andrew Murray, and Adam Stuckert for comments on early versions of the manuscript. This work was funded by the Explorers’ Club 2014 Youth Activity Grant, the Harvard Museum of Comparative Zoology’s Myvanwy M. and George M. Dick Scholarship Fund for Science Students, and a grant from the Harvard College Research Program to JRM, the Myvanwy M. and George M. Dick Scholarship Fund for Science Students and the Harvard College Research Program to SNC, and a Bauer Fellowship from Harvard University, the L’Oreal For Women in Science Fellowship, and the William F. Milton Fund from Harvard Medical School to LAO. EET and LAC acknowledges the support of Wikiri and the Saint Louis Zoo.

## Supplementary Figures

**Supplementary Figure 1.**
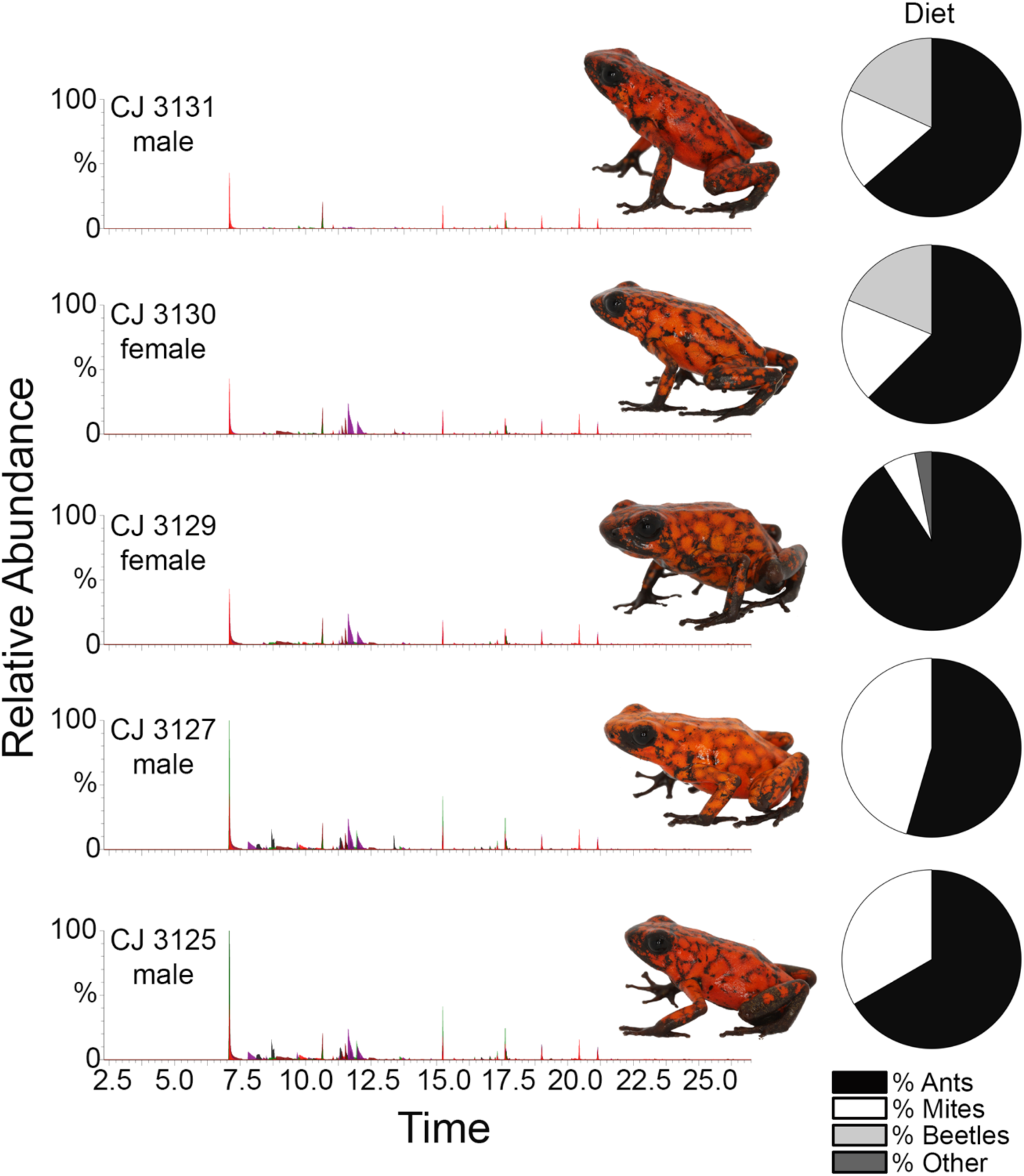
Individual variation in alkaloid profiles and morphology in the Little Devil frog *(Oophaga sylvatica)* Felfa population. (a) Skin alkaloid toxin profiles assayed by gas chromatography mass spectrometry (GCMS) show little individual variation in chemical defenses where each peak is a chemical and peak amplitude represents abundance. A picture of each individual is displayed in the right of each chromatogram. The pie chart on the right shows each frog's stomach content composition of percent ants (black), mites (white), beetles (light grey) or other arthropods (dark grey) by number.^100^ CJ 3097 male

**Supplementary Figure 2.**
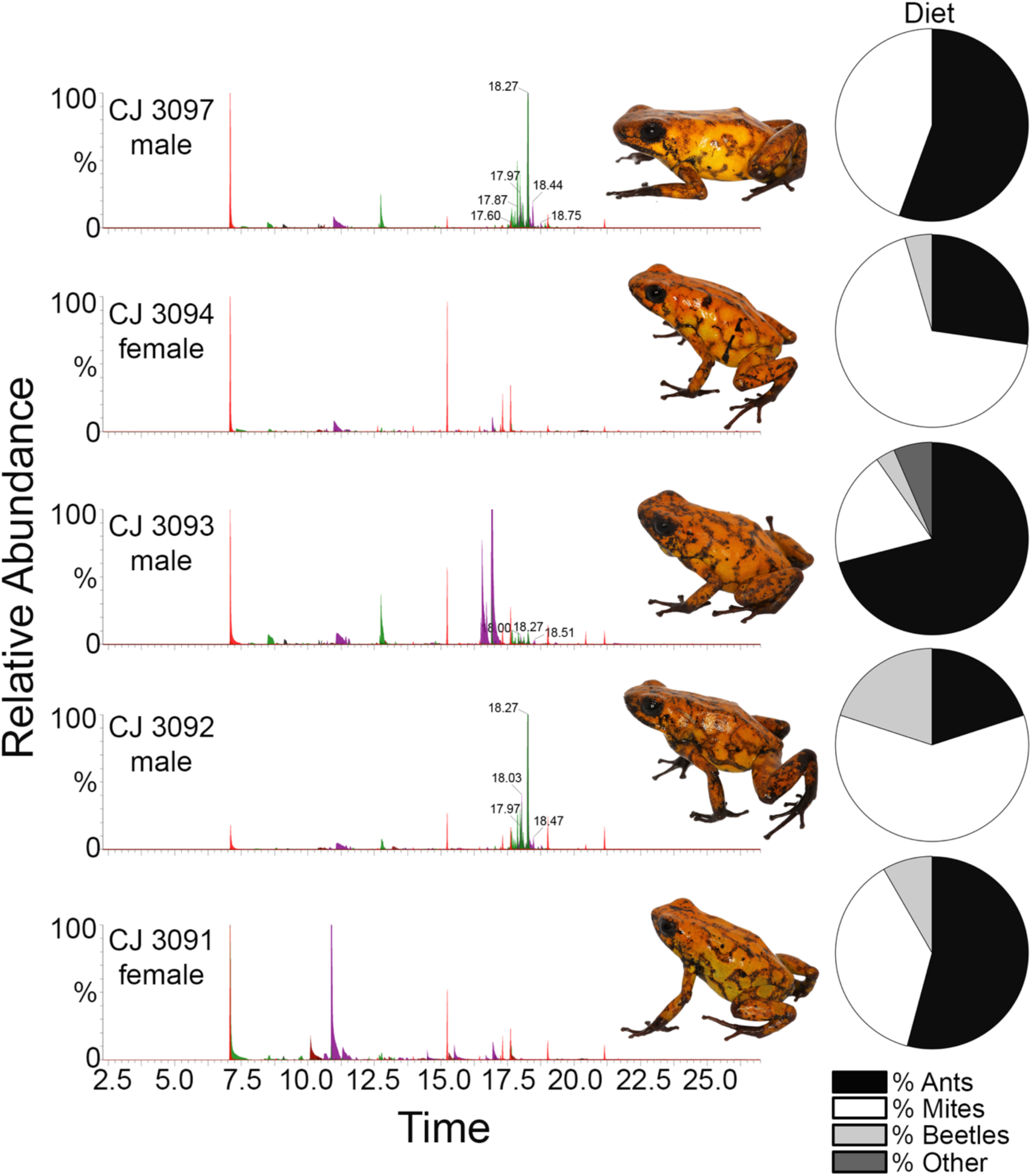
Individual variation in alkaloid profiles and morphology in the Little Devil frog *(Oophaga sylvatica)* Cristóbal Colón population. (a) Skin alkaloid toxin profiles assayed by gas chromatography mass spectrometry (GCMS) show individual variation in chemical defenses where each peak is a chemical and peak amplitude represents abundance. A picture of each individual is displayed in the right of each chromatogram. The pie chart on the right shows each frog's stomach content composition of percent ants (black), mites (white), beetles (light grey) or other arthropods (dark grey) by number.

**Supplementary Figure 3.**
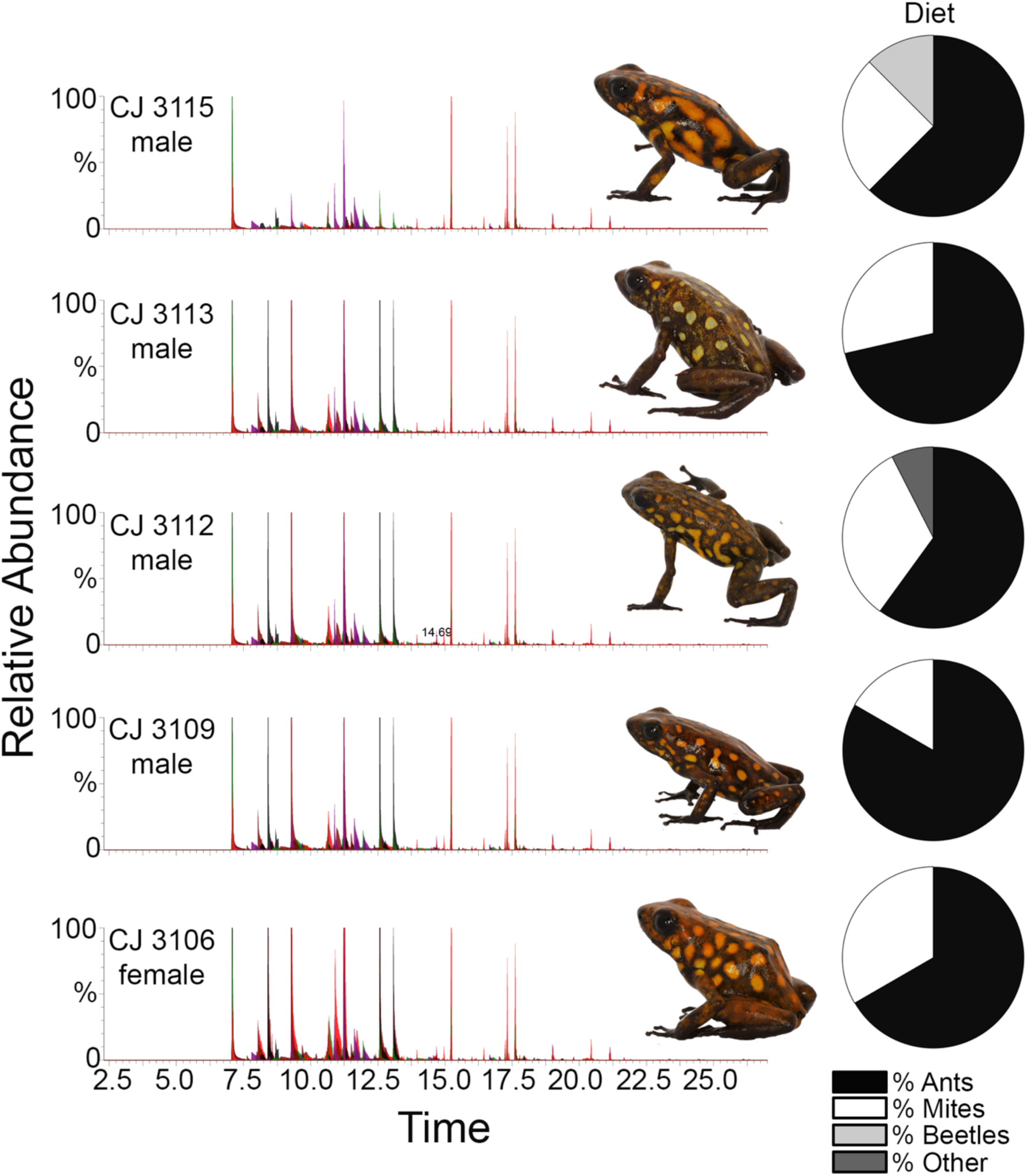
Individual variation in alkaloid profiles and morphology in the Little Devil frog *(Oophaga sylvatica)* Simón Bolívar population. (a) Skin alkaloid toxin profiles assayed by gas chromatography mass spectrometry (GCMS) show little individual variation in chemical defenses where each peak is a chemical and peak amplitude represents abundance. A picture of each individual is displayed in the right of each chromatogram. The pie chart on the right shows each frog's stomach content composition of percent ants (black), mites (white), beetles (light grey) or other arthropods (dark grey) by number.929 Supplementary Figure 4

**Supplementary Figure 4.**
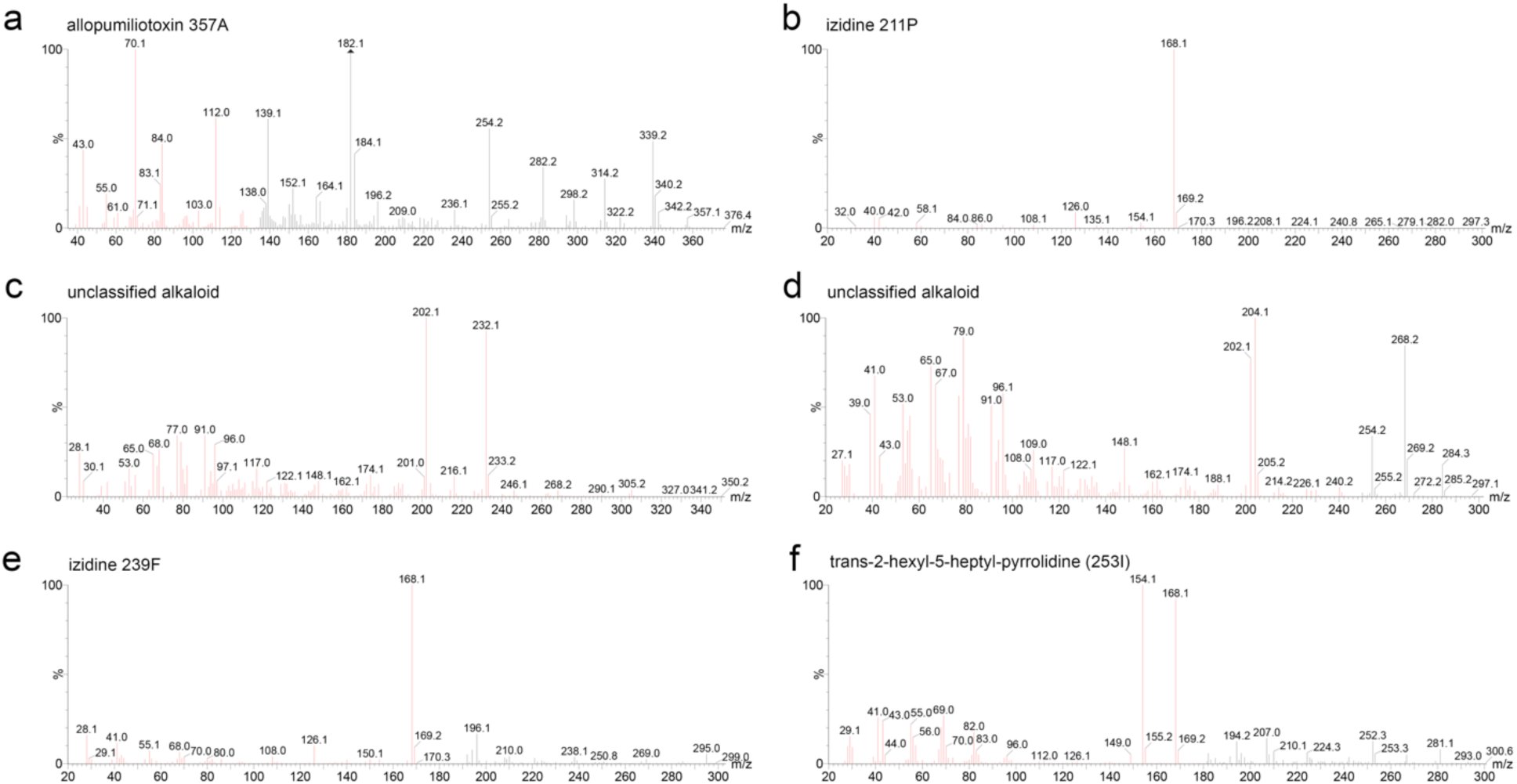
Ion mass spectra analysis of various features shown in Figure 2.

**Supplementary Figure 5.**
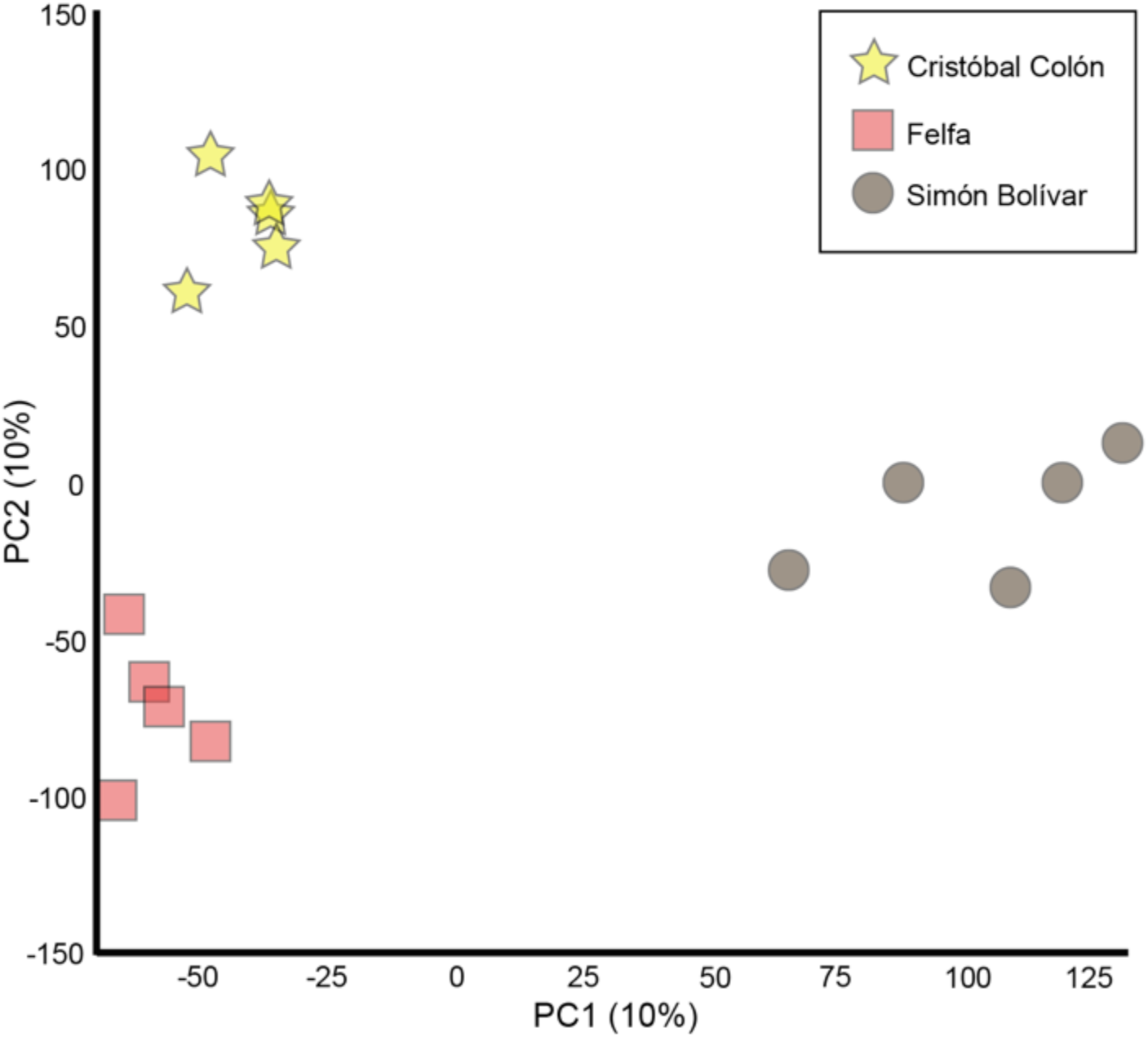
Principal component analysis of frog skin alkaloids based on liquid chromatography mass spectrometry analysis.

**Supplementary Figure 6.**
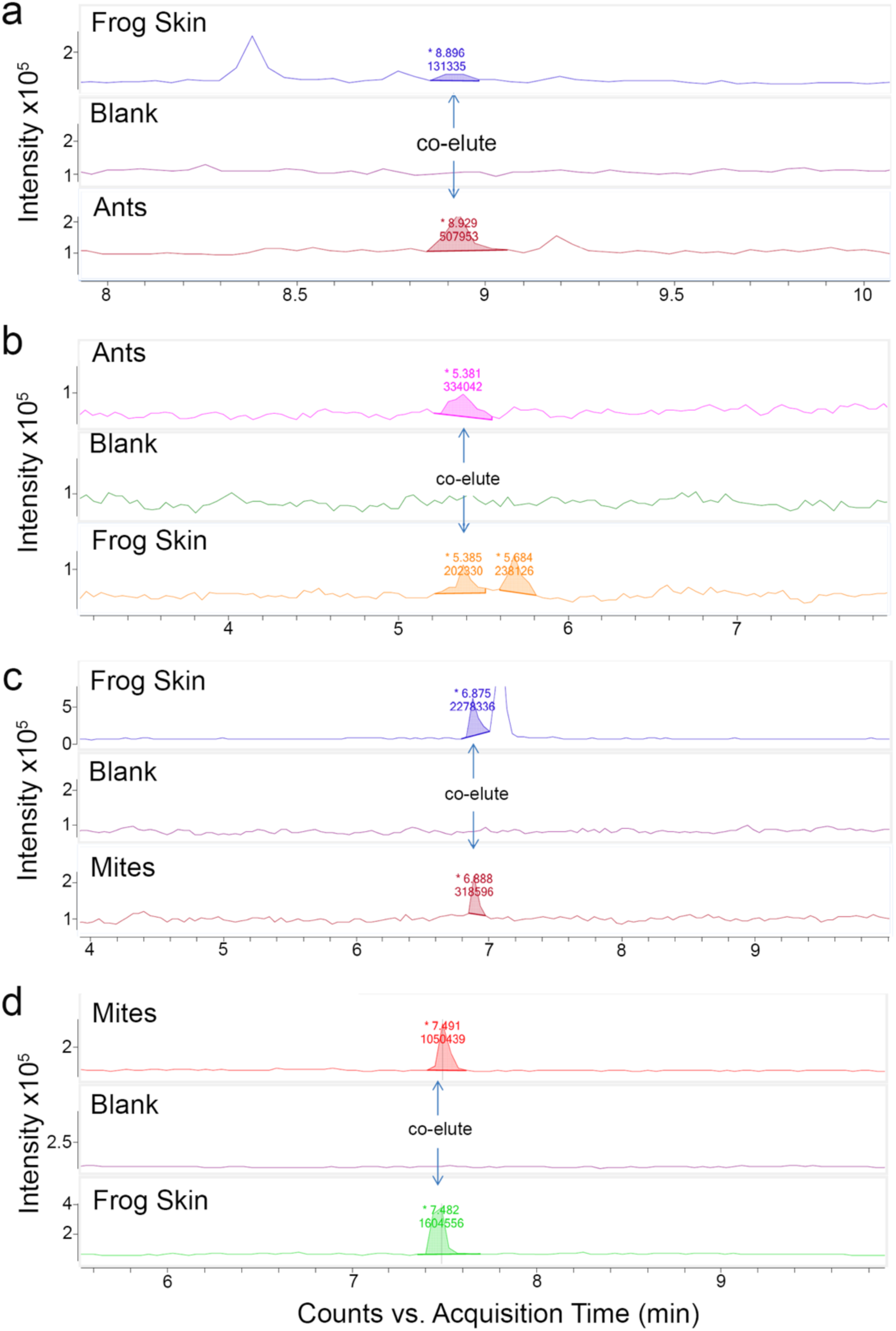
(M+H)^+^ product scans to compare retention times of frog and arthropod compounds and confirm co-elution. A methanol blank sample was run in between each frog skin and arthropod sample to ensure the matching compound was not due to carryover. (a) The major common response of the m/z 278 compound in the frog skin and ant sample (Figure 5a). (b) The major common response of the m/z 204 compound in the frog skin and ant sample (Figure 5b). (c) The major common response of the m/z 224 compound in the frog skin and mites sample (Figure 6a). (d) The major common response of the m/z 292 compound in the frog skin and mites sample (Figure 6b).

## Supplementary Tables

**Supplementary Table 1.** Ant identification using cytochrome oxidase 1. Assigned taxonomic identity by Sub-Family (Tribe) is listed in the first column and is based on the similarity to other ant cytochrome oxidase 1 sequences in the NCBI nr database. The sample ID indicates the voucher specimen number of the frog from whose stomach the ant was isolated (first four digits) and a three digit number assigned to prey items in the order they were retrieved from the stomach contents. The GenBank Accession number for each sequence is listed in the third column. The population of the frog from which each ant was isolated is listed in the fourth column. The closest arthropod hit for each specimen and its GenBank accession number is listed along with the similarity (percent identity) between the specimen and the closest GenBank match. The E-value statistic for the match is listed in the last column.

| Sub-Family<br>(Tribe) | Sample ID | GenBank<br>Accession | Frog<br>Population | BLASTn Species<br>(closest match) | GenBank<br>(closest match) | Similarity | E<br>Value |
| --- | --- | --- | --- | --- | --- | --- | --- |
| Myrmicinae<br>(Attini) | CJ 3111-001 | KU128492 | Simón Bolívar | *Octostruma* sp. | KF371477 | 94% | 0 |
| Myrmicinae<br>(Attini) | CJ 3111-002 | KU128470 | Simón Bolívar | *Wasmannia*<br>*aeropunctata* | KC985492 | 97% | 0 |
| Myrmicinae<br>(Attini) | CJ 3111-009 | KU128471 | Simón Bolívar | *Wasmannia*<br>*aeropunctata* | KC985492 | 96% | 0 |
| Myrmicinae<br>(Attini) | CJ 3111-013 | KU128473 | Simón Bolívar | *Wasmannia*<br>*aeropunctata* | KC985492 | 97% | 0 |
| Myrmicinae<br>(Attini) | CJ 3114-001 | KU128490 | Simón Bolívar | *Sericomyrmex* sp. | KC419855 | 94% | 0 |
| Myrmicinae<br>(Attini) | CJ 3116-001 | KU128487 | Simón Bolívar | *Cyphomyrmex* sp. | DQ353380 | 86% | 0 |
| Myrmicinae<br>(Attini) | CJ 3116-002 | KU128488 | Simón Bolívar | *Cyphomyrmex*<br>*minutus* | JQ617495 | 91% | 0 |
| Myrmicinae<br>(Attini) | CJ 3116-003 | KU128474 | Simón Bolívar | *Wasmannia*<br>*aeropunctata* | KC985492 | 97% | 0 |
| Myrmicinae<br>(Solenopsidini) | CJ 3116-006 | KU128483 | Simón Bolívar | *Solenopsis* sp. | KM224746 | 98% | 0 |
| Myrmicinae<br>(Solenopsidini) | CJ 3116-007 | KU128484 | Simón Bolívar | *Solenopsis* sp. | KM224746 | 98% | 0 |
| Myrmicinae<br>(Attini) | CJ 3116-012 | KU128478 | Simón Bolívar | *Wasmannia*<br>*aeropunctata* | KC985492 | 97% | 0 |
| Myrmicinae<br>(Attini) | CJ 3116-013 | KU128468 | Simón Bolívar | *Wasmannia*<br>*aeropunctata* | KC985492 | 97% | 0 |
| Myrmicinae<br>(Attini) | CJ 3116-016 | KU128472 | Simón Bolívar | *Wasmannia*<br>*aeropunctata* | KC985492 | 97% | 0 |
| Myrmicinae<br>(Attini) | CJ 3116-017 | KU128475 | Simón Bolívar | *Wasmannia*<br>*aeropunctata* | KC985492 | 97% | 0 |
| Myrmicinae<br>(Attini) | CJ 3116-022 | KU128469 | Simón Bolívar | *Wasmannia*<br>*aeropunctata* | KC985492 | 97% | 0 |
| Myrmicinae<br>(Solenopsidini) | CJ 3089-001 | KU128458 | Cristóbal<br>Colón | *Solenopsis* sp. | KC418686 | 85% | 0 |
| Myrmicinae<br>(Attini) | CJ 3089-006 | KU128496 | Cristóbal<br>Colón | *Pheidole* sp. | EF518348 | 93% | 0 |
| Myrmicinae<br>(Attini) | CJ 3089-008 | KU128459 | Cristóbal<br>Colón | *Solenopsis* sp. | KC418686 | 85% | 0 |
| Ectatomminae<br>(Ectatommini) | CJ 3089-009 | KU128479 | Cristóbal<br>Colón | *Gnamptogenys* sp. | KF371095 | 95% | 0 |
| Myrmicinae<br>(Attini) | CJ 3089-010 | KU128493 | Cristóbal<br>Colón | *Pheidole* sp. | KC418007 | 93% | 0 |
| Ectatomminae<br>(Ectatommini) | CJ 3089-014 | KU128480 | Cristóbal<br>Colón | *Gnamptogenys* sp. | KF371095 | 95% | 0 |
| Ectatomminae<br>(Ectatommini) | CJ 3089-016 | KU128481 | Cristóbal<br>Colón | *Gnamptogenys* sp. | KF371095 | 95% | 0 |
| Ectatomminae<br>(Ectatommini) | CJ 3089-017 | KU128482 | Cristóbal<br>Colón | *Gnamptogenys* sp. | KF371095 | 95% | 0 |
| Myrmicinae<br>(Attini) | CJ 3089-020 | KU128476 | Cristóbal<br>Colón | *Wasmannia*<br>*aeropunctata* | KC985492 | 97% | 0 |
| Myrmicinae<br>(Attini) | CJ 3095-010 | KU128489 | Cristóbal<br>Colón | *Cyphomyrmex* sp. | KC419123 | 91% | 0 |
| Myrmicinae<br>(Attini) | CJ 3096-017 | KU128463 | Cristóbal<br>Colón | *Wasmannia*<br>*aeropunctata* | KC985492 | 97% | 0 |
| Myrmicinae<br>(Attini) | CJ 3124-001 | KU128494 | Felfa | *Pheidole*<br>*glomericeps* | KM224728 | 98% | 0 |
| Myrmicinae<br>(Attini) | CJ 3124-007 | KU128495 | Felfa | *Pheidole*<br>*glomericeps* | KM224728 | 98% | 0 |
| Myrmicinae<br>(Attini) | CJ 3126-009 | KU128477 | Felfa | *Wasmannia*<br>*aeropunctata* | KC985492 | 96% | 0 |
| Myrmicinae<br>(Solenopsidini) | CJ 3128-001 | KU128485 | Felfa | *Solenopsis* sp. | KC418778 | 90% | 0 |
| Myrmicinae<br>(Solenopsidini) | CJ 3128-002 | KU128486 | Felfa | *Solenopsis* sp. | KC417942 | 85% | 0 |
| Myrmicinae<br>(Solenopsidini) | CJ 3128-003 | KU128491 | Felfa | *Solenopsis* sp. | KM224748 | 99% | 0 |
| Myrmicinae<br>(Attini) | CJ 3128-007 | KU128464 | Felfa | *Wasmannia*<br>*auropunctata* | KC985492 | 97% | 0 |
| Myrmicinae<br>(Attini) | CJ 3128-008 | KU128467 | Felfa | *Wasmannia*<br>*auropunctata* | KC985492 | 96% | 0 |
| Myrmicinae<br>(Attini) | CJ 3128-010 | KU128460 | Felfa | *Wasmannia*<br>*auropunctata* | KC985492 | 97% | 0 |
| Myrmicinae<br>(Attini) | CJ 3128-011 | KU128461 | Felfa | *Wasmannia*<br>*auropunctata* | KC985492 | 97% | 0 |
| Myrmicinae<br>(Attini) | CJ 3128-012 | KU128462 | Felfa | *Wasmannia*<br>*auropunctata* | KC985492 | 97% | 0 |
| Myrmicinae<br>(Attini) | CJ 3128-013 | KU128465 | Felfa | *Wasmannia*<br>*auropunctata* | KC985492 | 97% | 0 |
| Myrmicinae<br>(Attini) | CJ 3128-014 | KU128466 | Felfa | *Wasmannia*<br>*auropunctata* | KC985492 | 97% | 0 |
| Myrmicinae<br>(Crematogastrini) | CJ 3128-015 | KU128497 | Felfa | *Crematogaster sp.* | KC419078 | 94% | 0 |
| Myrmicinae<br>(Crematogastrini) | CJ 3132-002 | KU128456 | Felfa | *Crematogaster sp.* | KC417504 | 86% | 0 |
| Myrmicinae<br>(Crematogastrini) | CJ 3132-004 | KU128453 | Felfa | *Crematogaster sp.* | KC417504 | 86% | 0 |
| Myrmicinae<br>(Crematogastrini) | CJ 3132-005 | KU128457 | Felfa | *Crematogaster sp.* | KC417504 | 86% | 0 |
| Myrmicinae<br>(Crematogastrini) | CJ 3132-006 | KU128455 | Felfa | *Crematogaster sp.* | KC417504 | 86% | 0 |
| Myrmicinae<br>(Crematogastrini) | CJ 3132-007 | KU128454 | Felfa | *Crematogaster sp.* | KC417504 | 86% | 0 |

**Supplementary Table 2.** Mite identification using cytochrome 963 oxidase 1. Assigned taxonomic identity by Order is listed in the first column and is based on the similarity to other mite cytochrome oxidase 1 sequences in the NCBI nr database. The sample ID indicates the voucher specimen number of the frog from whose stomach the mite was isolated (first four digits) and a three digit number assigned to prey items in the order they were retrieved from the stomach contents. The GenBank Accession number for each sequence is listed in the third column. The population of the frog from which each mite was isolated is listed in the fourth column. The closest arthropod hit for each specimen and its GenBank accession number is listed along with the similarity (percent identity) between the specimen and the closest GenBank match. The E-value statistic for the match is listed in the last column.

| Order | Sample ID | GenBank Accession | Frog Population | BLASTn Species (closest match) | GenBank (closest match) | Similarity | E Value |
| --- | --- | --- | --- | --- | --- | --- | --- |
| Oribatida | CJ 3095-005 | KT947982 | Cristóbal Colón | *Archegozetes longisetosus* | HQ711372 | 99% | 0 |
| Oribatida | CJ 3095-007 | KT947987 | Cristóbal Colón | *Ceratozetidae* | KM831519 | 79% | 1e-96 |
| Oribatida | CJ 3096-002 | KT947981 | Cristóbal Colón | *Sarcoptiformes* | KM834143 | 84% | 4e-119 |
| Oribatida | CJ 3096-003 | KT947986 | Cristóbal Colón | *Berniniella hauseri* | KF293512 | 84% | 0 |
| Oribatida | CJ 3110-005 | KT947985 | Simón Bolívar | *Oppiella nova* | KM293453 | 78% | 2e-117 |
| Oribatida | CJ 3111-029 | KT947984 | Simón Bolívar | *Eremaeidae* | KM826715 | 79% | 2e-114 |
| Oribatida | CJ 3116-027 | KT947988 | Simón Bolívar | *Eremaeidae* | KM827121 | 79% | 7e-113 |
| Oribatida | CJ 3116-033 | KT947980 | Simón Bolívar | *Oppiella nova* | KF293453 | 79% | 4e-120 |
| Oribatida | CJ 3126-010 | KT947983 | Felfa | *Eueremaeus tetrosus* | KM839082 | 84% | 1e-142 |

